# PRS-LDmix: ancestry-mixed linkage disequilibrium reference panels for polygenic risk prediction in diverse and admixed cohorts

**DOI:** 10.64898/2026.11.24.767901

**Authors:** Grace Koo, Donghyung Lee

## Abstract

**Summary:** Polygenic risk scores (PRS) are less accurate outside European ancestry, which risks widening health disparities as they move into clinical use. One reason is linkage disequilibrium (LD), the correlation between nearby variants, which differs between ancestries. Bayesian methods such as PRS-CS model LD across all variants jointly, using a reference panel matched to the ancestry of the genome-wide association study (GWAS). Multi-ancestry extensions such as PRS-CSx pair each ancestry with its own panel and require separate summary statistics for each ancestry. However, these are often incomplete or restricted, so scores are frequently built from the combined meta-analysis, whose LD is a mixture that no single panel matches. In practice, the majority-ancestry panel is typically used. Here, we introduce PRS-LDmix, which builds ancestry-specific LD panels from a 32,953-individual, 29-population reference, aligned to shared blocks and SNPs, and averages them in the cohort’s ancestry proportions, replacing PRS-CS’s LD input without changing its prior or sampler. In simulated GWAS mixing European with African, East Asian or South Asian ancestry, the mixed panel matched the true LD more closely than the majority-ancestry panel. Once the ancestries were balanced, the choice of panel shifted the scores beyond PRS-CS’s own run-to-run variation, most for European and African ancestry, and PRS-LDmix was more accurate than the majority-ancestry panel, though not always more accurate than the other ancestry’s own panel.

**Availability and implementation:** PRS-LDmix is implemented as an R package with a C++ backend and Python scripts that extend PRS-CS. It is freely available under the GPL (≥2) at https://github.com/statsleelab/PRS-LDmix.

## 1 Introduction

Polygenic risk scores (PRS) summarize genome-wide genetic effects into a single predictor of complex trait liability and disease risk (Lewis and Vassos, 2020; Torkamani et al., 2018). For individual *i*, the score is a weighted sum over single nucleotide polymorphisms (SNPs),

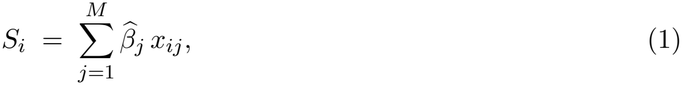

where *x*_*ij*_ ∈ {0, 1, 2} counts the copies of SNP *j*’s effect allele carried by individual *i* and 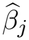 is that allele’s estimated effect. The weights are estimated from genome-wide association study (GWAS) summary statistics in a discovery sample, and the scores are computed and evaluated in a separate target sample. By quantifying an individual’s genetic predisposition to a disease, PRS may help identify people who would benefit from earlier screening or prevention (Torkamani et al., 2018).

A central challenge in computing PRS is linkage disequilibrium (LD), the non-random association of alleles at different loci. Nearby SNPs are rarely separated by recombination, so their genotypes are correlated. Because a GWAS tests each SNP on its own, a SNP’s estimated effect captures not only its own causal effect but also those of the SNPs in LD with it. Summing these marginal estimates double-counts shared signal, which can distort risk estimates and reduce predictive accuracy. Every PRS method must therefore account for LD (Choi et al., 2020; Vilhjálmsson et al., 2015).

The traditional way to handle LD is clumping and thresholding (C+T) (Chang et al., 2015; Choi and O’Reilly, 2019). Clumping retains the most significant SNP in each genomic region and removes neighbouring SNPs whose LD with it exceeds a threshold. Thresholding then keeps only SNPs whose GWAS *p*-value falls below a cut-off. C+T avoids double counting, but it discards the information carried by the removed SNPs and does not shrink noisy estimates.

LD-aware Bayesian methods instead model all SNPs jointly. PRS-CS (Ge et al., 2019) and LDpred2 (Privé et al., 2020) combine GWAS summary statistics with an LD matrix to re-estimate each SNP’s effect, accounting for its correlation with its neighbours. PRS-CS partitions the genome into 1,703 approximately independent LD blocks, inferred from European LD with LDetect (Berisa and Pickrell, 2016), and places a continuous shrinkage prior on SNP effects that shrinks weak signals strongly while leaving large effects largely intact. LDpred2 combines the summary statistics, local LD and a prior on effect sizes in a Bayesian model. Because individual-level GWAS data are rarely shared, both methods estimate LD from an external reference panel, such as the 1000 Genomes Project (The 1000 Genomes Project Consortium, 2015) or UK Biobank (Bycroft et al., 2018) panels distributed with PRS-CS, and take a single panel chosen to match the ancestry of the GWAS.

That choice matters because LD differs across populations. Differences in demographic history and allele frequencies give each ancestry group its own LD structure. PRS methods therefore need a reference panel matched to the ancestry of the GWAS. LD differences are also one reason PRS lose accuracy across populations (Lewis and Vassos, 2020; Wang et al., 2020). This exacerbates a broader problem: most GWAS participants are of European ancestry, and PRS are substantially less accurate in other populations (Martin et al., 2019). Multi-ancestry methods such as PRS-CSx (Ruan et al., 2022) model ancestry-specific GWAS jointly, each paired with its own ancestry-matched panel, but they require the per-ancestry summary statistics. These are often incomplete or restricted: some ancestry groups are withheld or too small, and some contributing cohorts do not permit genome-wide release of their results (Mahajan et al., 2022; Meng et al., 2024). The combined meta-analysis is then the largest available result, and its LD is a mixture that no single ancestry’s panel matches. In practice, the majority ancestry’s panel is typically used: the Global Biobank Meta-analysis Initiative, for example, used a European panel for its multi-ancestry meta-analyses and suggests the dominant ancestry’s panel for convenience, while noting that a panel mirroring the ancestry composition of the discovery GWAS is ideal when in-sample LD is unavailable (Wang et al., 2023).

PRS-LDmix addresses this case by matching the LD panel to the cohort. It builds ancestry-specific LD panels from the 33KG reference panel (Lee and Bacanu, 2024) on shared blocks and SNPs, and mixes them inside PRS-CS in the cohort’s ancestry proportions. The mixed panel replaces PRS-CS’s LD input, but its prior and Gibbs sampler are unchanged. In simulated mixed-ancestry GWAS, the mixed panel was closer to the true LD of the GWAS than the majority-ancestry panel. The choice of panel mattered more as the two ancestries approached equal proportions, shifting the scores by more than PRS-CS’s own run-to-run variation. In these more balanced cohorts, PRS-LDmix was also more accurate than the majority-ancestry panel, though not always as accurate as the other ancestry’s own panel.

Combining reference LD across ancestries to match a cohort has precedent in other summary-statistics analyses. DISTMIX (Lee et al., 2015) imputes association statistics at unmeasured SNPs in mixed-ethnicity cohorts, using a reference panel weighted by estimated or user-specified ethnic proportions. Adapt-Mix (Park et al., 2015) combines all available reference panels to estimate local LD in arbitrary populations, learning the mixture proportions from the summary statistics by how well they impute observed *z*-scores, and was evaluated for imputation and joint testing. Both describe the LD of a single pooled sample of the ancestries, which includes a between-ancestry term and is computed from reference genotypes. By contrast, PRS-LDmix averages finished, block-wise LD panels in the format PRS-CS reads. This average is the LD of a GWAS meta-analysed across ancestries and needs no allele frequencies (Supplementary Section S1). PRS-LDmix is also built for polygenic prediction, for which neither earlier method was evaluated.

Because scores are often built from multi-ancestry meta-analysed summary statistics, matching the LD panel to the cohort, rather than defaulting to its majority ancestry, is a simple change to how they are used, yet once the ancestries are balanced it shifts the scores systematically, beyond PRS-CS’s own run-to-run variation.

## 2 Methods

### 2.1 Building ancestry LD panels

#### Reference panel

PRS-LDmix estimates LD from the 33KG reference panel (Lee and Bacanu, 2024), whose 32,953 individuals come from 29 populations grouped into five superpopulations: African (AFR; 817 individuals), admixed American (AMR; 533), East Asian (ASN; 10,800), European (EUR; 20,281) and South Asian (SAS; 522). We estimate each superpopulation’s LD from its constituent reference samples as the covariance of a mixture of its subpopulations, weighted by their sample shares, which includes within- and between-population terms (Lee et al., 2015, Section 2.1); this is the estimator implemented in GAUSS (Lee and Bacanu, 2024). SNPs are filtered to minor allele frequency ≥ 0.01 within each superpopulation, and each block’s correlation matrix 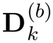 is corrected to be positive definite with an eigenvalue floor.

#### Shared blocks

PRS-CS treats LD as block-diagonal. Mixing panels block by block requires every ancestry’s panel to partition each chromosome identically, so PRS-LDmix defines blocks once per chromosome and applies them to every ancestry (Fig. 1). We use fixed 250-kb tiles spanning the base-pair range of the reference index. They are simple and depend on no ancestry’s LD structure, so they treat every ancestry alike, and they are used throughout this paper. Truncating LD at a fixed distance, rather than at LD-inferred boundaries, is common in summary-statistics PRS methods; for example, LDpred and LDpred2 ignore correlations beyond a fixed window around each variant (Privé et al., 2020; Vilhjálmsson et al., 2015). PRS-LDmix can also build every panel on a second layout, EUR-derived cut points: the block boundaries inferred from PRS-CS’s published EUR panel (Ge et al., 2019), which follow the European LD blocks that PRS-CS uses (Berisa and Pickrell, 2016). Users can therefore keep PRS-CS’s own block structure when mixing panels. Panels can be restricted to HapMap3 SNPs (The International HapMap 3 Consortium, 2010), as PRS-CS’s own panels are.

**Figure 1:**
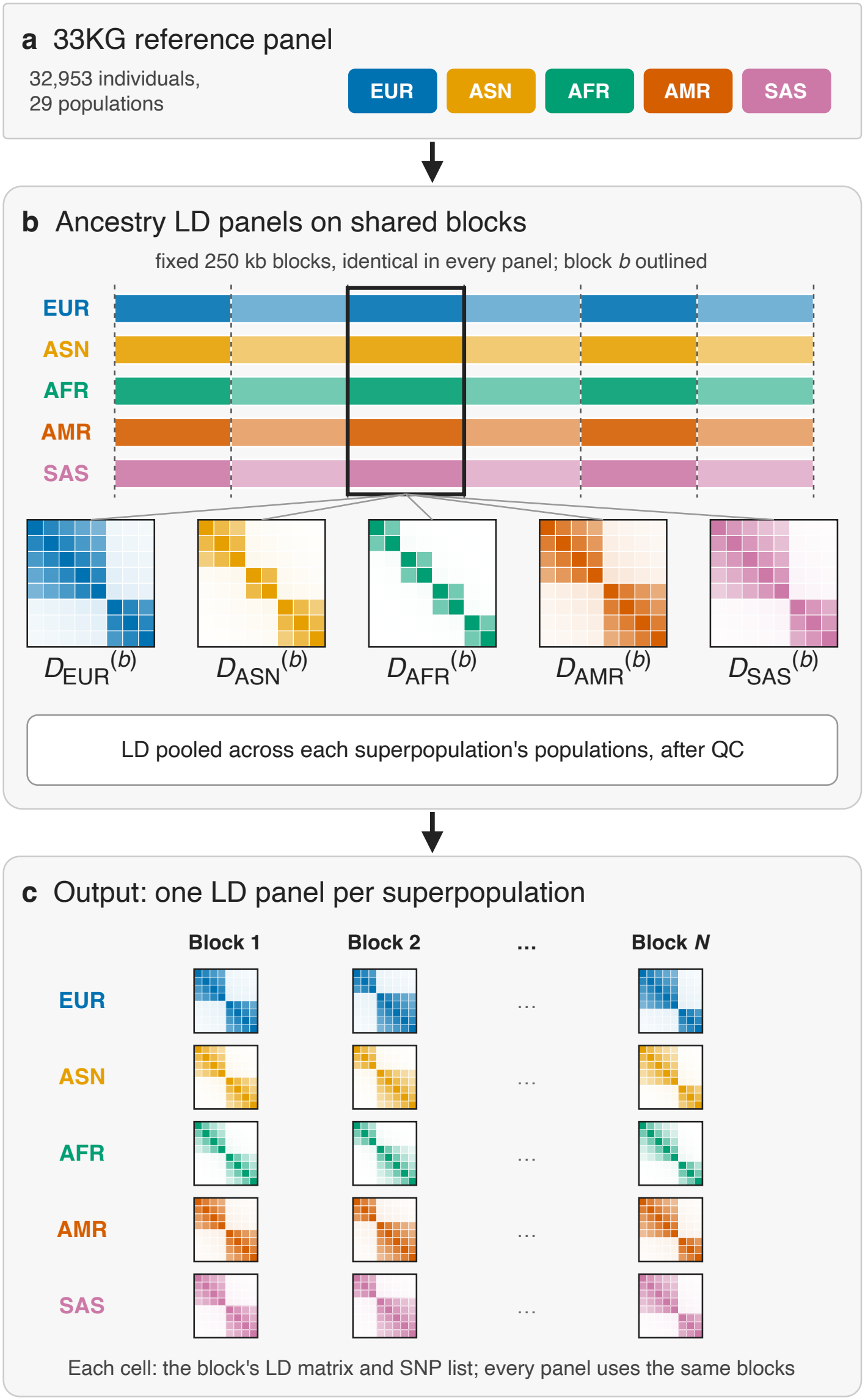
Schematic of how PRS-LDmix builds ancestry LD panels. (a) The 33KG reference panel groups its populations into five superpopulations (AFR, AMR, ASN, EUR and SAS). (b) LD is estimated for each superpopulation on the same fixed 250-kb blocks and (c) written as one PRS-CS-format panel per superpopulation. LD matrices are illustrative.

### 2.2 Mixing panels for a cohort

#### SNP alignment

Panels are mixed entry by entry, so within each block they must describe the same SNPs, in the same order, with the same effect allele (Fig. 2). All panels use the common genomic blocks of Section 2.1 and are built from the same reference index, so each SNP carries the same alleles in every panel. Because filtering on allele frequency can keep different SNPs for different ancestries, PRS-LDmix restricts each block to the SNPs present in every panel and in the GWAS summary statistics, orders them identically across panels, and aligns every panel to the GWAS effect allele with the same allele flips before mixing (Supplementary Section S2).

**Figure 2:**
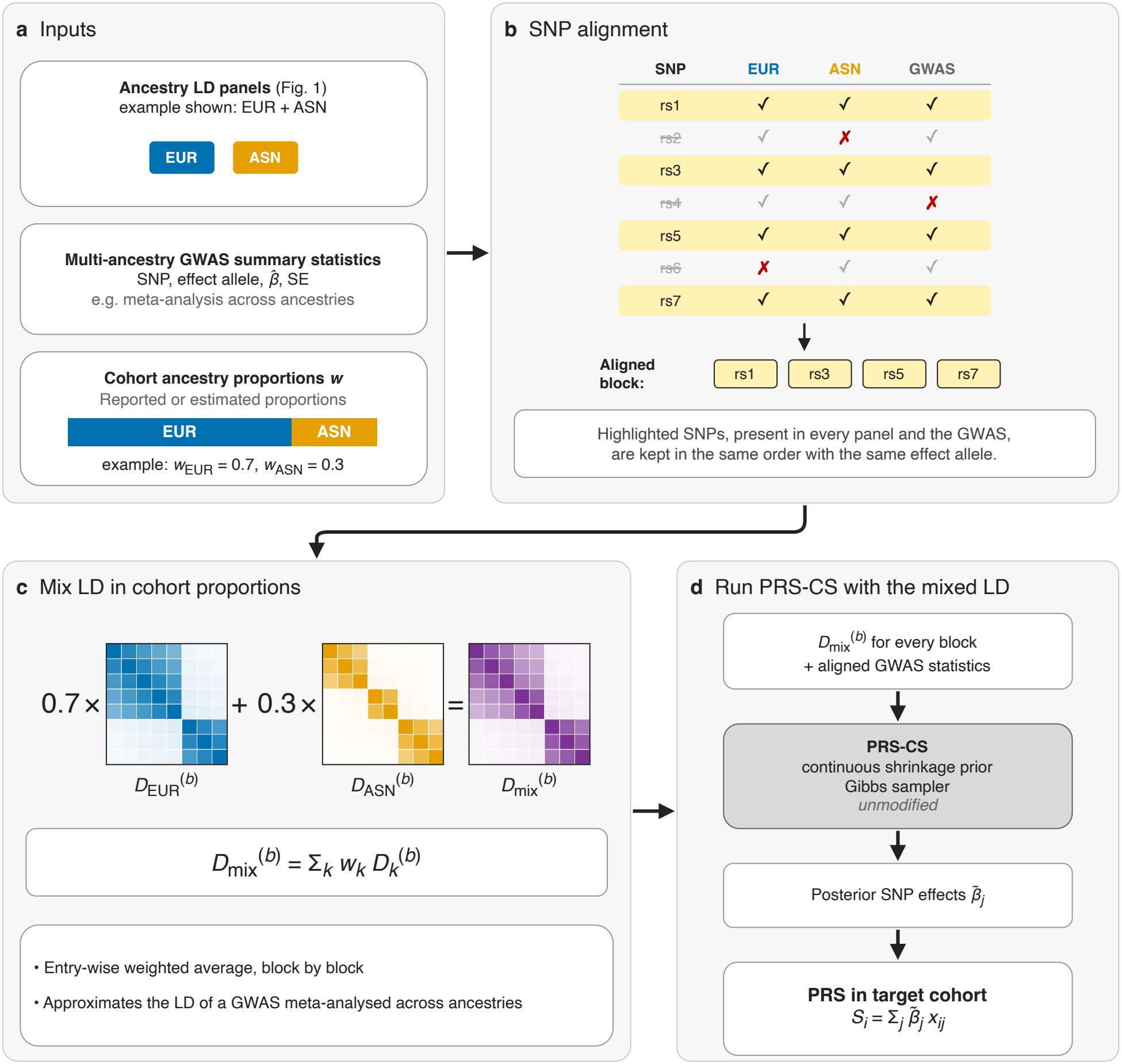
Schematic of how a user runs PRS-LDmix, shown for a cohort that is 70% EUR and 30% ASN. (a) The inputs are ancestry LD panels (Fig. 1), GWAS summary statistics and the cohort’s ancestry proportions *w*; (b) each block is restricted to the SNPs shared by every panel and the GWAS, in a common order and allele coding; (c) block LD matrices are averaged with weights *w* (Equation (2)); and (d) the mixed matrices replace PRS-CS’s LD input, leaving its prior and Gibbs sampler unchanged. LD matrices are illustrative.

#### Mixed LD matrices

PRS-LDmix constructs

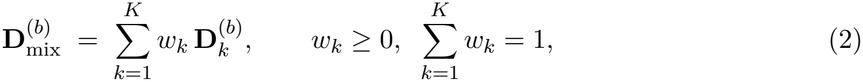

where 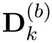 is the reference LD correlation matrix of superpopulation *k* in block *b*, and *K* is the number of superpopulations mixed. The mixing proportions are supplied by the user and should match the GWAS cohort’s ancestry composition, taken from its reported composition or estimated from the data. If independent ancestry-specific GWAS are combined by sample-size-weighted Stouffer meta-analysis (Willer et al., 2010), with *w*_*k*_ the sample shares, the same mixture of the discovery samples’ LD matrices is the null covariance of the combined *z*-statistics. PRS-LDmix uses reference-panel estimates to approximate this covariance. Applicability to other GWAS designs, and the further assumptions needed for the non-null mean model, are discussed in Supplementary Section S1.

Each panel’s block matrices are made positive definite with a small eigenvalue floor, so 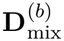, as a convex combination of them, is positive definite as well, although it can be ill-conditioned. When panels are built, the same shrinkage towards the identity, 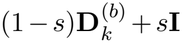 with *s* ∈ [0, 1), can optionally be applied to every panel; because the weights sum to one, the mixture of shrunk panels is then 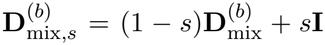, whose eigenvalues are at least *s*. The analyses reported here use no shrinkage (*s* = 0). PRS-LDmix substitutes the mixed matrix for each block’s LD matrix in PRS-CS and leaves its prior and Gibbs sampler unchanged. It has two components: the R function `generate_prscs_ld_panels()` builds the aligned single-ancestry panels in PRS-CS’s HDF5 layout, and the Python script `PRScs_mix.py` mixes them block by block and runs PRS-CS, taking the same options as PRS-CS plus a list of panel directories and matching mixing proportions. The script runs alongside an unmodified PRS-CS installation, from which it imports the sampler. Installation and usage are documented in the PRS-LDmix repository (https://github.com/statsleelab/PRS-LDmix).

### 2.3 Simulation design

We evaluated PRS-LDmix in simulations on chromosome 22, using the real genotypes of the 33KG panel and simulating only the phenotypes and GWAS summary statistics. We studied three ancestry pairs (EUR+ASN, EUR+SAS and EUR+AFR), each at three GWAS compositions (90:10, 70:30 and 50:50, the EUR share first). Every population was split at random into reference (50%), discovery (25%) and target (25%) individuals. The reference individuals are used only to build the LD panels, with the fixed 250-kb blocks, as a public reference panel would be. The discovery individuals are the GWAS participants, whose genotypes give the simulated GWAS its LD. The target individuals are held out and scored. Because the panels and the GWAS come from different individuals, the panels carry realistic estimation error. We build the simulated summary statistics in four steps: a phenotype model in each ancestry group, a GWAS in each group, a meta-analysis of the groups, and a nominal sample size that sets the strength of the signal.

#### Notation

The analysis SNPs, *m* of them, are the panels’ SNPs with no missing genotype in any discovery or target set and with variation in every discovery set. For ancestry group *k*, let **X**_*k*_ be the *n*_*k*_ × *m* matrix of discovery genotypes: one row per discovery individual, one column per SNP, and each entry the number of copies (0, 1 or 2) of that SNP’s effect allele. We standardize each SNP within the discovery set, subtracting its mean and dividing by its standard deviation (denominator *n*_*k*_), and call the result **Z**_*k*_; every column of **Z**_*k*_ then has mean zero and variance one. The LD matrix of the discovery individuals is 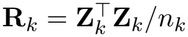. It is an *m* × *m* matrix whose entry for SNPs *j* and *l* is the correlation between their genotypes in group *k*, with ones on the diagonal. The composition weight *w*_*k*_ is group *k*’s share of the GWAS (*w*_*k*_ *>* 0, ∑_*k*_ *w*_*k*_ = 1); at 70:30, for example, *w*_EUR_ = 0.7. The LD of the combined GWAS is then **R**_mix_ = ∑_*k*_ *w*_*k*_**R**_*k*_, each group’s LD weighted by its share. This is the quantity that Equation (2) estimates from reference panels, so the simulation gives PRS-LDmix a known target.

#### Step 1: phenotype model

Within each group, every discovery individual is given a phenotype,

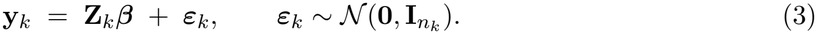

The vector ***β*** holds the true effect of each SNP, measured per standard deviation of genotype. The same effects apply in every ancestry group, so differences between groups come only from their genotypes and LD; the causal effects of common variants are largely shared across ancestries (Wang et al., 2020). Multiplying the genotypes by the effects, **Z**_*k*_***β***, gives each individual’s genetic value: the part of the phenotype due to the SNPs. The noise ***ε***_*k*_ is an independent standard normal draw for each individual and stands for the environment and everything else; it has variance one, which sets the scale for the strength of the genetic signal. In the null simulation, every effect is zero, so the phenotype is pure noise and unrelated to genotype; in the non-null simulation, a small fraction of SNPs have non-zero effects (see True effects below).

#### Step 2: GWAS within each group

A GWAS tests each SNP on its own for association with the phenotype. Because the genotypes are standardized and the noise has variance one, the *z*-statistics of all SNPs are, to a close approximation for small effects,

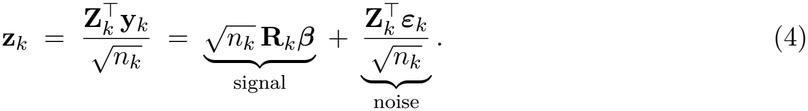

The first expression is the usual association statistic: for each SNP, the sum over individuals of genotype times phenotype, scaled by the sample size. Substituting the phenotype model of Equation (3) splits it into two parts. The signal part, 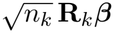, is what the true effects contribute. Because it is multiplied by the LD matrix, each SNP’s signal is the sum of the effects of all SNPs it is correlated with: a SNP with no effect of its own still shows an association if it is in LD with a causal SNP. This is the double counting that PRS methods must undo, and why they need the LD matrix. The signal also grows with 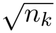, so larger GWAS detect effects more strongly. The noise part comes from the random ***ε***_*k*_. Given the genotypes, it is normally distributed with mean zero and covariance **R**_*k*_: chance associations are correlated between SNPs in exactly the way their genotypes are, so neighbouring SNPs tend to look associated together.

#### Step 3: meta-analysis across groups

The group-level GWAS are combined by sample-size-weighted Stouffer meta-analysis, 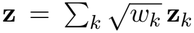: each group’s *z*-statistics are weighted by the square root of its share of the total sample and added. This is the meta-analysis design for which Equation (2) gives the LD (Supplementary Section S1). Its noise part is

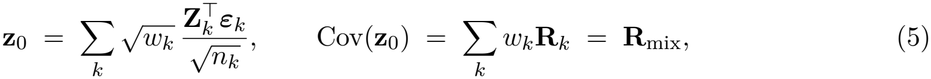

given the discovery genotypes. The first expression adds up each group’s noise from Equation (4) with the meta-analysis weights. The second says how that noise is correlated between SNPs. Each group’s noise has covariance **R**_*k*_; the groups’ noise is independent, so the covariances add; and squaring the weight 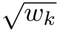 gives *w*_*k*_. The combined noise is therefore correlated according to **R**_mix_, the composition-weighted blend of the groups’ LD. This is the central point of the design: the LD of a meta-analysed multi-ancestry GWAS is a mixture of the ancestries’ LD, which is what PRS-LDmix approximates. In the null simulation there is no signal, so **z** = **z**_0_: pure noise, with the LD of the combined GWAS.

#### Step 4: nominal sample size

The discovery groups are small, from about a hundred to a few thousand individuals (Table 2), which would give too weak a signal for a realistic GWAS. We therefore compute the signal part as if group *k* had *w*_*k*_*N* individuals, with a nominal total sample size *N* = 200,000: in the signal term of Equation (4), 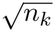 is replaced by 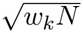 . The noise part **z**_0_ still comes from the actual discovery genotypes, because its distribution, with covariance **R**_*k*_ in each group, does not depend on the sample size. Combining the groups as in Step 3 gives

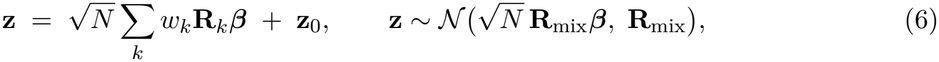

since each group’s signal is multiplied first by its meta-analysis weight 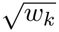 and then by 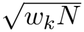, and 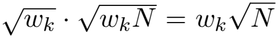 The first expression is the *z*-statistic of the simulated GWAS: signal plus the noise of Equation (5). The second gives its distribution. On average, the *z*-statistics equal the true effects spread through the combined LD and scaled by 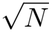, and around that average they vary as correlated noise with the combined LD. This is what a meta-analysed GWAS of *N* individuals in these proportions would look like, with the LD of the discovery individuals.

#### True effects

In each non-null replicate, 1% of the analysis SNPs are chosen as causal, uniformly at random without replacement. Their effects are drawn independently from N (0, 1), and all other SNPs have effect zero. All effects are then multiplied by one constant so that ***β***^*⊤*^**R**_mix_***β*** = ∑_*k*_ *w*_*k*_∥**Z**_*k*_***β***∥^2^*/n*_*k*_ = *v*_*g*_, with *v*_*g*_ = 0.01. In words, ∥**Z**_*k*_***β***∥^2^*/n*_*k*_ is the variance of the genetic values among group *k*’s discovery individuals, so *v*_*g*_ is the variance of the genetic values over the combined GWAS, measured against noise variance one. Rescaling keeps the signal strength the same in every replicate, while the causal SNPs, their effects and the noise are drawn anew. These sparse normal effects are not the continuous shrinkage prior that PRS-CS assumes; the mismatch is the same for every LD panel compared.

#### Input to PRS-CS

The summary statistics were passed to PRS-CS as effect estimates 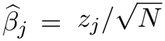 with standard errors 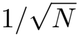 and `–n_gwas` = *N*, from which PRS-CS recovers 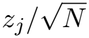. Because **z** is computed from the discovery genotypes, it carries their LD in full, including LD across block boundaries, which every panel ignores, and PRS-LDmix is evaluated with estimated panels rather than with the generating covariance **R**_mix_ itself.

#### Analyses and scoring

In every setting, each non-null replicate was analysed with PRS-CS (*ϕ* auto, default sampler settings) using the EUR panel, the second ancestry’s panel and the PRS-LDmix panel at the generating composition; each single-panel analysis was also rerun with a new sampler seed (Section 2.4). PRS-CS returned effects on the standardized scale. Target genotypes **X**_*T*_ of each ancestry were standardized with that ancestry’s discovery means and standard deviations, giving **Z**_*T*_ ; each analysis’s scores are 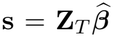, and the true genetic values are **g** = **Z**_*T*_ ***β***. Each setting has 100 replicates. The null model of Equation (5) defines the true LD compared in Section 3.2. Table 1 summarises the null and non-null simulations, and Table 2 lists the settings of each ancestry pair.

**Table 1:**
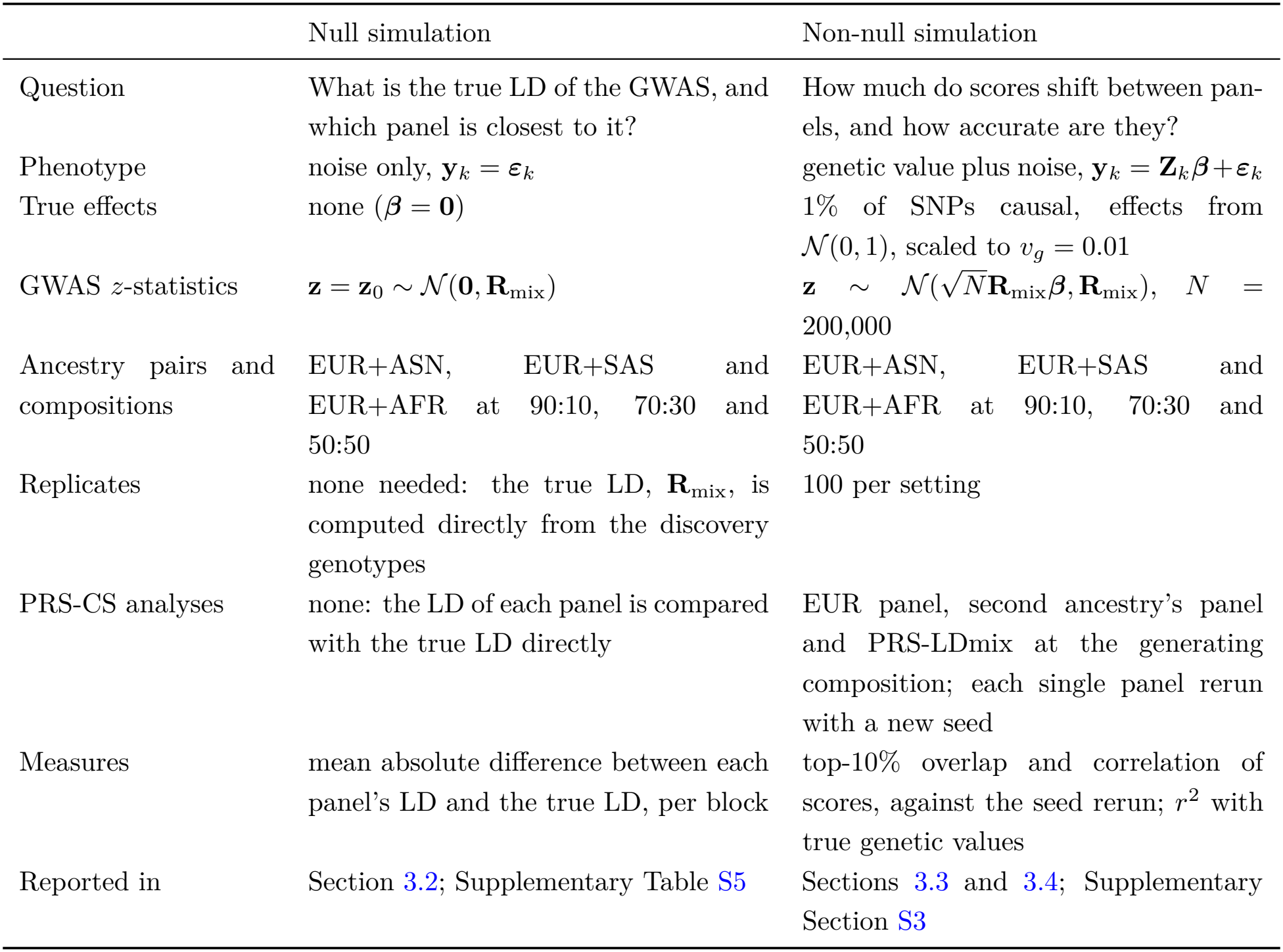
The null and non-null simulations at a glance (Section 2.3). Both use chromosome 22, the reference/discovery/target split, LD panels with fixed 250-kb blocks and Stouffer meta-analysis of the ancestry groups. Further settings are in Table 2.

|  | Null simulation | Non-null simulation |
|---|---|---|
| Question | What is the true LD of the GWAS, and which panel is closest to it? | How much do scores shift between pan-els, and how accurate are they? |
| Phenotype | noise only, $\mathbf{y}_k = \boldsymbol{\varepsilon}_k$ | genetic value plus noise, $\mathbf{y}_k = \mathbf{Z}_k\boldsymbol{\beta} + \boldsymbol{\varepsilon}_k$ |
| True effects | none ($\boldsymbol{\beta} = \mathbf{0}$) | 1% of SNPs causal, effects from $\mathcal{N}(0, 1)$, scaled to $v_g = 0.01$ |
| GWAS *z*-statistics | $\mathbf{z} = \mathbf{z}_0 \sim \mathcal{N}(\mathbf{0}, \mathbf{R}_{\text{mix}})$ | $\mathbf{z} \sim \mathcal{N}(\sqrt{N}\mathbf{R}_{\text{mix}}\boldsymbol{\beta}, \mathbf{R}_{\text{mix}})$, $N = 200{,}000$ |
| Ancestry pairs and compositions | EUR+ASN, EUR+SAS and EUR+AFR at 90:10, 70:30 and 50:50 | EUR+ASN, EUR+SAS and EUR+AFR at 90:10, 70:30 and 50:50 |
| Replicates | none needed: the true LD, $\mathbf{R}_{\text{mix}}$, is computed directly from the discovery genotypes | 100 per setting |
| PRS-CS analyses | none: the LD of each panel is compared with the true LD directly | EUR panel, second ancestry's panel and PRS-LDmix at the generating composition; each single panel rerun with a new seed |
| Measures | mean absolute difference between each panel's LD and the true LD, per block | top-10% overlap and correlation of scores, against the seed rerun; *r*<sup>2</sup> with true genetic values |
| Reported in | Section 3.2; Supplementary Table S5 | Sections 3.3 and 3.4; Supplementary Section S3 |

**Table 2:** Simulation and analysis settings of the ancestry-pair simulations (Section 2.3).

| Item | Value |
|---|---|
| Ancestry pairs | EUR and ASN, EUR and AFR, EUR and SAS |
| GWAS ancestry mixes | 90:10, 70:30 and 50:50 for each pair |
| Chromosome | 22, HapMap3 SNPs |
| LD blocks | fixed 250 kb, shared by both panels |
| Simulated GWAS | 1% causal SNPs with effects shared across ancestries; nominal *N* = 200,000 |
| Analysis SNPs per pair | EUR and ASN 14,233 (142 causal); EUR and AFR 15,693 (157 causal); EUR and SAS 15,997 (160 causal) |
| Signal variance | *v<sub>g</sub>* = 0.01 (in units of residual variance) |
| GWAS (discovery) individuals | EUR 5,070, ASN 2,701, AFR 205, SAS 132 |
| Scored (target) individuals | EUR 5,070, ASN 2,699, AFR 204, SAS 129 |
| LD panels | reference individuals only; no additional convex shrinkage; eigenvalue floor 10<sup>−8</sup> applied for numerical stability |
| Replicates | 100 per setting |
| PRS-LDmix blend | each panel weighted by its ancestry’s share of the GWAS |
| PRS-CS | *ϕ* auto, sampler defaults (1,000 iterations, 500 burn-in, thinning 5) |
| Sampler seeds | 1000 + replicate; rerun 2000 + replicate |

### 2.4 Metrics and controls

Score agreement between two analyses is measured by top-tail overlap, the share of target individuals in one analysis’s top score decile who are also in the other’s top decile, and by the correlation between their scores. Accuracy is measured by *r*^2^ between scores and true genetic values. Comparisons are paired within replicate, since every analysis uses the same simulated GWAS and scores the same target individuals.

Each replicate also reruns each single-panel analysis with a different sampler seed. This seed rerun measures the run-to-run variation of PRS-CS itself, and every score comparison is made against it: PRS-LDmix’s agreement with a single panel is compared with the agreement between that panel and its own seed rerun, so a change of panel matters only if it moves scores further than a new seed does. The main reference analysis is the EUR panel, the panel standard PRS-CS would use for these EUR-majority cohorts; comparisons with the second ancestry’s panel are in the supplement. Results are means over replicates with Monte Carlo standard errors (the standard deviation over replicates divided by the square root of their number); paired differences use the standard error of the per-replicate difference. The layout pilot and convergence checks report 95% paired *t* intervals.

## 3 Results

### 3.1 Panels reproduce PRS-CS’s LD

Single-ancestry PRS-LDmix panels reproduce the LD of PRS-CS’s published panels, and with a single panel PRS-LDmix reproduces PRS-CS’s output exactly. We checked the two components separately. Genome-wide panels (chromosomes 1–22, HapMap3 SNPs) built for all five superpopulations, on both the fixed 250-kb blocks and the EUR-derived cut points, passed every structural and matrix check on all 22 chromosomes: each panel lists the same SNPs and alleles in the same order, and every block’s LD matrix is symmetric and positive definite, with a unit diagonal. Their within-block correlations agreed with those of PRS-CS’s published 1000 Genomes panel for the matching population, compared on SNP pairs in the same block in both panels; the median correlation over chromosomes ranged from 0.987 to 0.999 across superpopulations with fixed blocks and from 0.964 to 0.997 with EUR-derived cut points (Supplementary Table S6). The 33KG panel includes the 1000 Genomes samples, so this confirms that the panels are built correctly rather than that they replicate in independent samples. Given a single panel, `PRScs_mix.py` wrote posterior effect files byte-identical to those of unmodified `PRScs.py` with the same inputs and seed, in 80 of 80 files (Supplementary Table S7).

### 3.2 A mixed cohort’s LD is closer to the mixed panel than the majority panel

To see whether the LD panel differs systematically from the LD of a mixed-ancestry GWAS, we used the null simulations to compare the true LD of the simulated GWAS with the LD each analysis assumes, at three GWAS compositions (90:10, 70:30 and 50:50). The true LD is **R**_mix_ in Equation (5), the discovery cohorts’ LD blended at the GWAS composition; it uses no PRS-CS output and is the same in every replicate. The analyses assume either the PRS-LDmix blend of the reference panels at the same composition or the EUR panel that standard PRS-CS would use. For each block, we measured the mean absolute difference from the true LD over all SNP pairs.

Fig. 3 illustrates this for one block of chromosome 22 (block 75, 34.55–34.80 Mb) in the EUR+AFR pair. As the two ancestries approach equal proportions, the EUR panel’s LD departs further from the true LD, while the PRS-LDmix blend stays much closer to it. Fig. 4 shows the same block for EUR+ASN, where the EUR panel departs less from the true LD (EUR+SAS is in Supplementary Fig. S1). This is expected, because EUR and ASN are less divergent than EUR and AFR. European and East Asian populations share an out-of-Africa bottleneck, whereas LD decays fastest with distance in African populations (The 1000 Genomes Project Consortium, 2015). Block 75 was chosen for its visible LD structure; by the EUR panel’s misfit it ranks 7–10 of 138 blocks for EUR+AFR and 16–27 of 137 for EUR+ASN across the three compositions, so it shows the pattern more clearly than a typical block.

**Figure 3:**
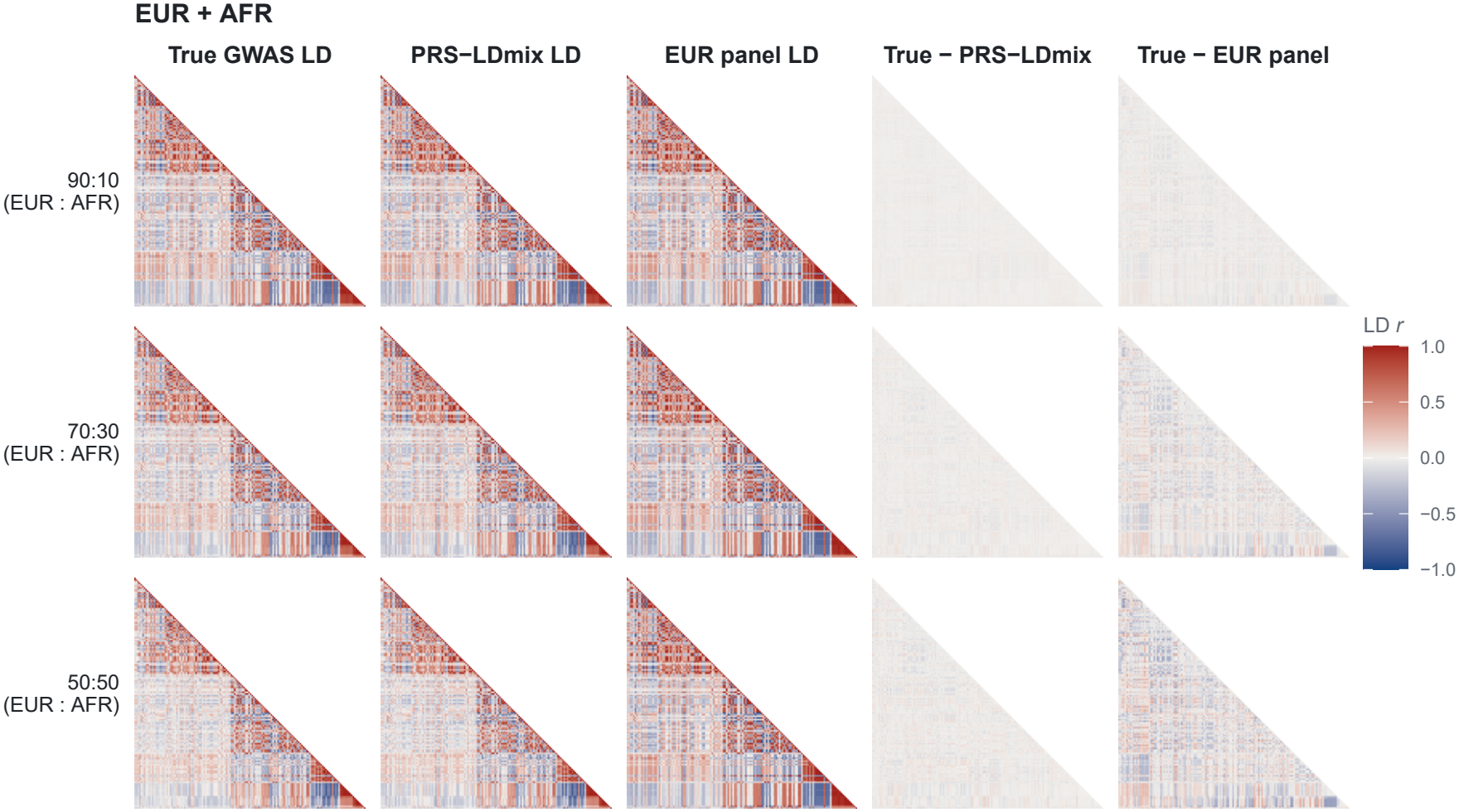
The LD of a simulated EUR+AFR GWAS and the LD each method assumes, in one 250-kb block on chromosome 22 (block 75, 34.55–34.80 Mb, 144 SNPs; Section 3.2). Rows are GWAS compositions (EUR : AFR); columns are the true GWAS LD, the PRS-LDmix blend of the reference panels, the EUR panel used by standard PRS-CS, and the differences of the last two from the true LD. One colour scale throughout; lower triangle only.

**Figure 4:**
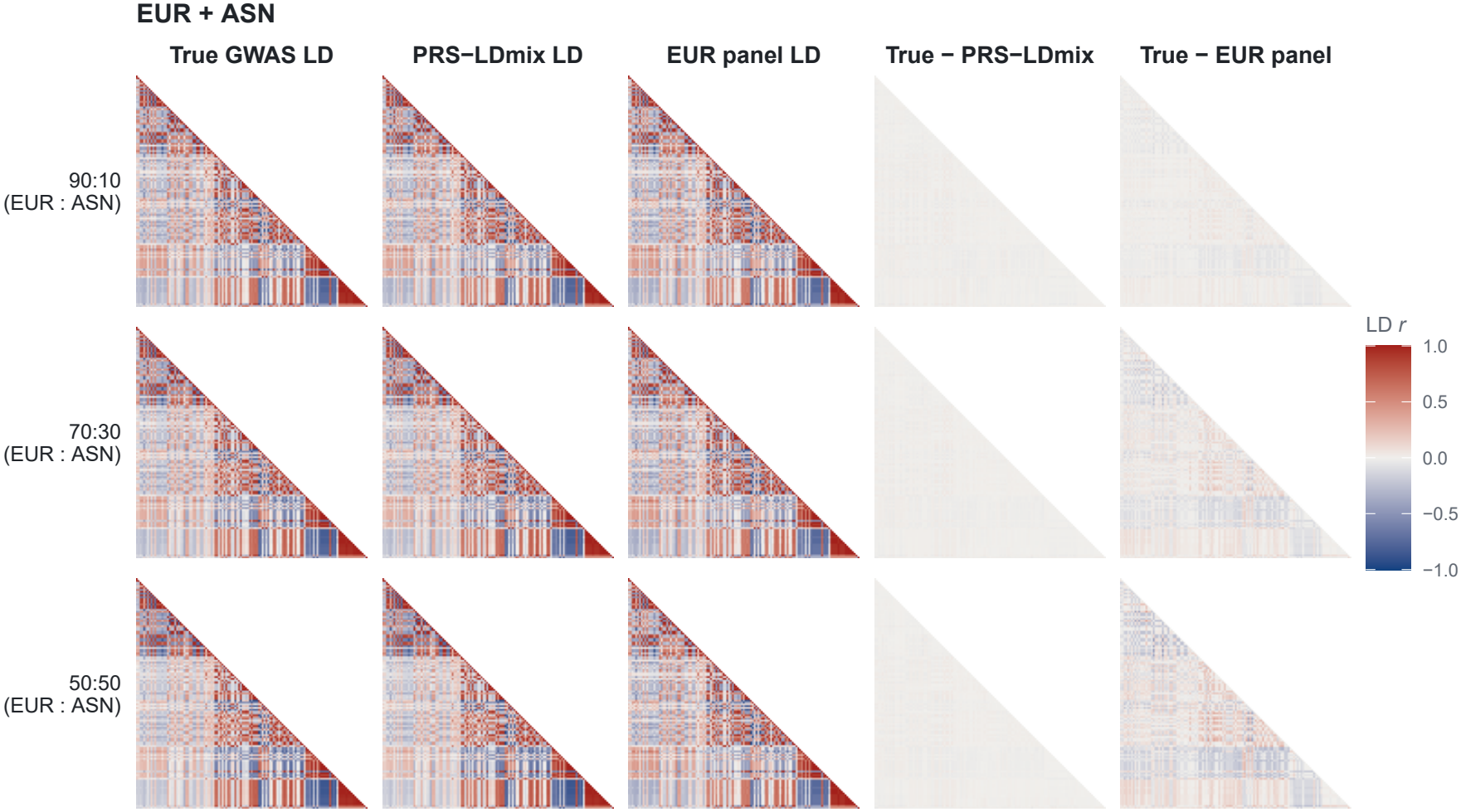
As Fig. 3, for EUR+ASN in the same block; rows are GWAS compositions (EUR : ASN). ASN is less divergent from EUR than AFR is, so the EUR panel departs less from the true LD than in Fig. 3.

The pattern holds across the chromosome. PRS-LDmix was closer to the true LD than the EUR panel in 138 of 138 blocks for EUR+AFR and 137 of 137 for EUR+ASN at every composition, and in at least 133 of 138 for EUR+SAS (blocks with at least two analysis SNPs; Supplementary Table S5). The EUR panel’s median per-block difference from the true LD rose from 0.020 at 90:10 to 0.078 at 50:50 for EUR+AFR; at 50:50 it was 0.063 for EUR+ASN and 0.052 for EUR+SAS, the pair closest to EUR. The median difference for PRS-LDmix stayed near 0.011 at every composition for EUR+ASN but rose to 0.033 for EUR+AFR and 0.039 for EUR+SAS at 50:50, so for EUR+SAS its advantage at 50:50 was smallest. The second ancestry’s own panel fitted no better than the EUR panel: at 50:50 its median difference was similar to the EUR panel’s for EUR+ASN and larger for EUR+AFR and EUR+SAS. The remaining difference for PRS-LDmix is the gap between the LD of the reference and discovery individuals of each ancestry, weighted by the composition. It grows with the weight on the small AFR and SAS samples (205 AFR and 132 SAS discovery individuals, against 2,701 ASN and 5,070 EUR), so it reflects sampling noise in small samples more than error in the blend.

### 3.3 Scores shift beyond seed noise, more with balance and divergence

Which LD panel is used shifts the scores systematically, by more than PRS-CS’s own run-to-run variation, and the shift grows as the two ancestries approach equal proportions. In each replicate, we compared the scores from PRS-LDmix at the generating composition with those from the EUR panel that standard PRS-CS would use, by the overlap of their top 10%. The reference point is the EUR panel rerun with a new sampler seed: its overlap with the original EUR run is the agreement expected from sampler randomness alone, so PRS-LDmix shifts the scores only where its overlap falls below the seed rerun’s. Both comparisons use the same simulated GWAS and score the same target individuals, so they are paired within replicate (Fig. 5). Values below are means over 100 replicates per setting; standard errors are in Supplementary Table S1.

**Figure 5:**
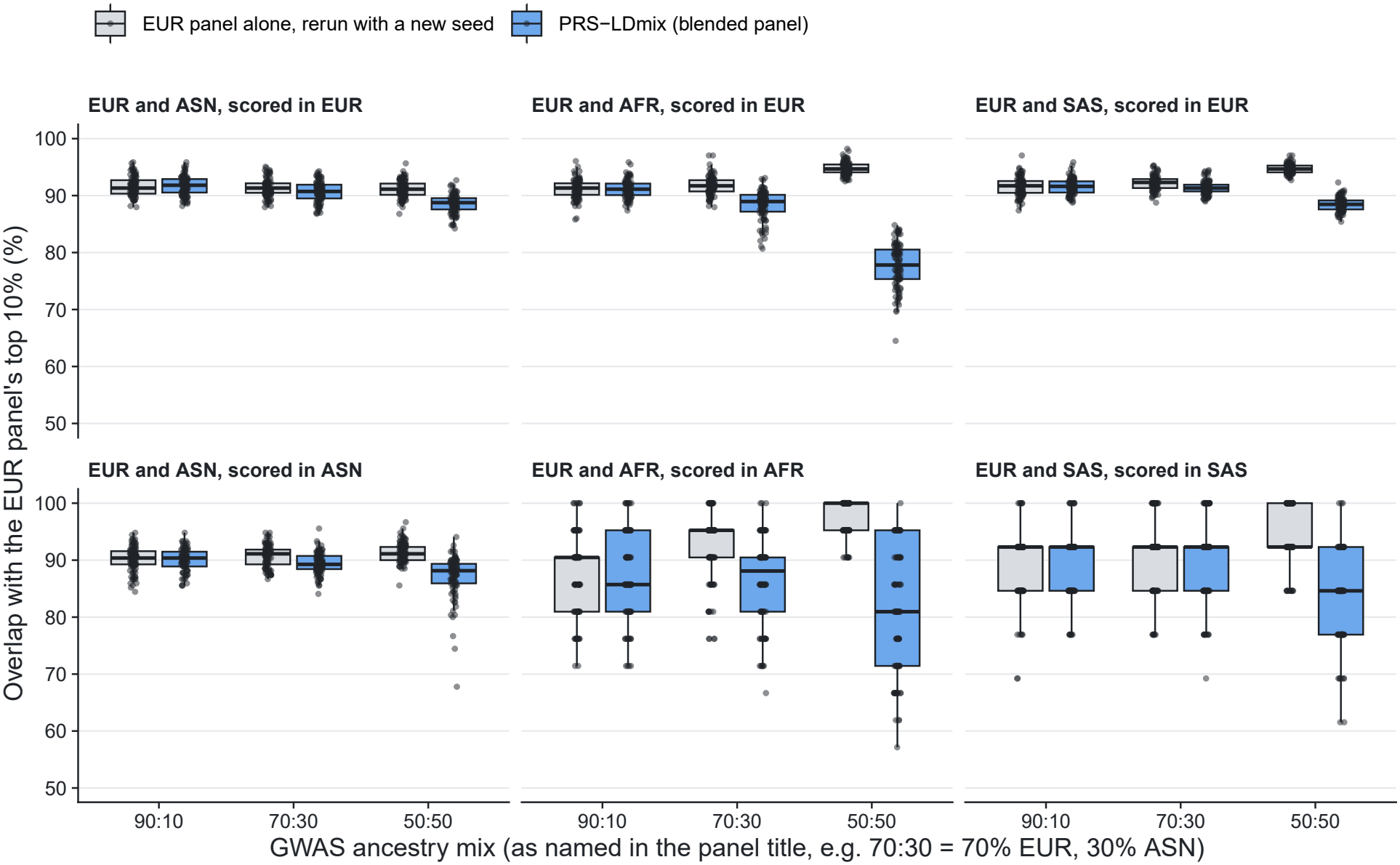
Agreement of the top 10% of scores with single-panel EUR PRS-CS, by ancestry pair (columns) and GWAS composition (horizontal axis), for target individuals from EUR (top row) or from the pair’s second ancestry (bottom row). Overlap is the share of an analysis’s top 10% that is also in the EUR panel’s top 10%; grey boxes are the EUR panel rerun with a new sampler seed (sampler randomness alone), and blue boxes are PRS-LDmix at the generating composition. Each point is one of 100 replicates (chromosome 22, fixed 250-kb blocks, *ϕ* auto).

At 90:10, PRS-LDmix and the EUR panel agreed as closely as two runs of the EUR panel did: in every pair and in both target ancestries, the two overlaps differed by at most 0.7 percentage points. As the compositions became more balanced, PRS-LDmix moved away from the EUR panel by more than a new seed does. The shift was largest for EUR+AFR, the most divergent pair, where at 50:50 the top-10% overlap with the EUR panel was 77.7%, against 94.7% for the seed rerun, in EUR target individuals, and 82.3% against 97.7% in AFR target individuals. For EUR+ASN at 50:50 it was 88.6% against 91.1% in EUR target individuals. Relative to its seed rerun, EUR+SAS shifted more than EUR+ASN at 50:50, although SAS is closer to EUR: PRS-LDmix’s overlap with the EUR panel was similar in the two pairs (88.5% and 88.6%), but the EUR panel’s seed rerun agreed more closely with the original run for EUR+SAS (94.7% against 91.1%). The AFR and SAS target groups are small, so their overlaps vary more between replicates (Fig. 5). The same pattern held for the top 5% and 20% and for the correlation of scores (Supplementary Tables S2 and S3), and on chromosome 2, which has several times as many SNPs as chromosome 22 (Supplementary Section S6.1).

### 3.4 Accuracy compared with single-ancestry panels

PRS-LDmix matched the majority-ancestry panel at 90:10 (*r*^2^ with the true genetic values within 0.003 in every pair) and was more accurate once the ancestries were balanced: for EUR+AFR at 50:50, *r*^2^ was 0.277 for PRS-LDmix against 0.202 for the EUR panel in EUR target individuals (Supplementary Table S4). For EUR+ASN at 50:50, PRS-LDmix was the most accurate of the three analyses in both target groups: *r*^2^ was 0.593 against 0.586 for the EUR panel and 0.579 for the ASN panel in EUR target individuals, and 0.593 against 0.579 and 0.581 in ASN target individuals. In the other two pairs, however, it was not always as accurate as the best single panel. The second ancestry’s own panel was more accurate at 50:50 for EUR+AFR (0.302 against 0.277 in EUR target individuals) and, by a smaller margin, for EUR+SAS (0.246 against 0.241), in target individuals of both ancestries; for EUR+AFR, the AFR panel was already ahead at 70:30 in AFR target individuals. The AFR panel’s advantage was also present on chromosome 2 (Supplementary Section S6.1), and it remained when every panel was built from the GWAS’s own discovery individuals, whose LD is the LD the GWAS was simulated under (Supplementary Table S10), so it is not caused by a mismatch between reference and discovery LD.

## 4 Discussion

PRS-LDmix provides LD reference panels matched to a cohort’s ancestry composition, built so that panels from different ancestries can be mixed block by block, and runs them in PRS-CS without changes to its inference. Its main use is not a gain in average accuracy. The choice of LD panel is itself a modelling decision that shifts individual scores systematically, beyond run-to-run variation, and PRS-LDmix makes that choice match the cohort rather than defaulting to one ancestry.

The simulations show when this choice matters. The LD of a mixed-ancestry GWAS is a blend of its ancestries’ LD, and the majority-ancestry panel departs further from it as the ancestries become more balanced (Section 3.2). The scores follow the same pattern. When one ancestry dominates (90:10), PRS-LDmix and the majority panel agree as closely as two runs of PRS-CS with different seeds, so defaulting to the majority panel costs little. As the cohort becomes more balanced, the two move apart by more than a new seed does, most for the most divergent pair, EUR+AFR (Section 3.3). Because every comparison is paired with a seed rerun, this shift reflects the LD assumed rather than sampler randomness. EUR+SAS shifted more relative to its seed rerun than its closeness to EUR would suggest, because sampler randomness moved its scores less, not because PRS-LDmix moved them more. The shift is not, in itself, a gain in accuracy: PRS-LDmix was clearly more accurate than the majority panel in balanced cohorts but not always more accurate than the other ancestry’s panel (Section 3.4).

PRS-LDmix can build every panel on either of two block layouts. Fixed 250-kb blocks, used throughout this paper, are simple and depend on no ancestry’s LD. EUR-derived cut points follow the block structure of PRS-CS’s published EUR panel, so users can keep PRS-CS’s own layout, and panels on both layouts passed the same checks (Section 3.1). In a layout pilot, the EUR-derived cut points and larger fixed blocks raised the accuracy of every method for EUR+ASN, with mixed effects for EUR+AFR (Supplementary Section S7). Because the cut points follow European LD, they are not neutral across ancestries in the way fixed blocks are.

### Limitations

The evaluation uses simulated traits, mainly on chromosome 22, with a check on chromosome 2. The mixing rule models a GWAS meta-analysed across ancestries; for a pooled GWAS, and especially one of admixed individuals, whose genomes carry additional LD from their mixed ancestry, it is an approximation, closest when the analysis adjusts for ancestry. The mixing proportions must be supplied by the user. The AFR and SAS samples are small, both in the reference panels and among the discovery individuals that define the simulated GWAS’s LD. PRS-CS approximates LD as block-diagonal, so LD across block boundaries is ignored for every panel. All analyses use PRS-CS’s default sampler, which retains about 100 posterior draws; longer chains raised the accuracy of every method by a similar amount and left the comparisons between panels essentially unchanged (Supplementary Section S6.3).

### Future work

The most important next step is to evaluate PRS-LDmix on real rather than simulated data. In simulation the true LD of the GWAS and the true genetic values are known; in real cohorts, the scores can still be compared against a seed rerun, and accuracy can be measured against observed phenotypes. We plan to use two or three multi-ancestry datasets, each with ancestry-specific GWAS summary statistics and an independent target sample of each ancestry with measured phenotypes. Within each dataset, meta-analysing GWAS run on subsamples of the discovery cohorts would give a range of compositions, as in the simulations. This would test whether the score shifts and the accuracy seen here hold when the GWAS LD, the genetic architecture and the ancestry labels are not simulated.

The mixed matrices are ordinary block-diagonal LD matrices, so they could also be supplied to other LD-based PRS methods, such as LDpred2 (Privé et al., 2020) and SBayesRC (Zheng et al., 2024), after conversion where a method needs its own LD format, to test whether the score shifts seen with PRS-CS carry over. In the longer term, we aim to release precomputed genome-wide panels and a service in which users choose a PRS method, supply their GWAS summary statistics, ancestry proportions and target genotypes, and receive scores built from mixed panels on the shared blocks. Because many cohorts cannot share individual-level genotypes, the service would also be offered as a container that runs locally.

## Acknowledgements

Computations for this work were run on the RedHawk high-performance computing cluster, supported by Research Computing Support at Miami University.

## Data availability

The 33KG reference panel (Lee and Bacanu, 2024) is available from the GAUSS package website (https://statsleelab.github.io/gauss/articles/ref_33KG.html). PRS-CS’s 1000 Genomes LD reference panels are available from the PRS-CS repository (https://github.com/getian107/PRScs). No individual-level data other than the reference panel were used. The code that builds the panels, simulates the GWAS summary statistics and computes the results, and the curated results used to build all tables and figures, are in the PRS-LDmix repository (https://github.com/statsleelab/PRS-LDmix). PRS-CS was run with the repository’s `PRScs_mix.py`, using the settings in Section 2.3.

## Supplementary Information

### Part A: Method details

#### S1 Derivation and scope of the mixing rule

This section shows why averaging the ancestries’ LD matrices, as in Equation (2), gives the LD of a meta-analysed multi-ancestry GWAS, and when that holds.

##### Result

Suppose *K* independent ancestry-specific GWAS, with *n*_*k*_ individuals in study *k* and *N* = ∑_*k*_ *n*_*k*_ in total, are combined by sample-size-weighted Stouffer meta-analysis (Willer et al., 2010), 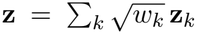 with *w*_*k*_ = *n*_*k*_*/N* . Let **R**_*k*_ be the LD correlation matrix of study *k*’s individuals and ***β***_*k*_ the true SNP effects in ancestry *k* on its standardized genotype scale. Then

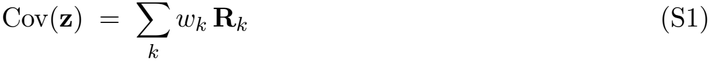

and the meta-analysed marginal effects 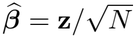 have mean

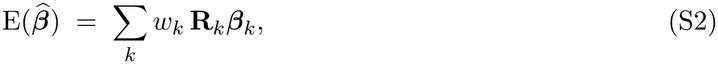

which equals **R**_mix_***β***, with **R**_mix_ = ∑_*k*_ *w*_*k*_**R**_*k*_, when the effects are shared, ***β***_*k*_ = ***β*** for every *k*.

**Proof**. Within study *k*, let **Z**_*k*_ be the *n*_*k*_ × *m* matrix of standardized genotypes, so that 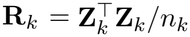, and let the phenotype be **y**_*k*_ = **Z**_*k*_***β***_*k*_ + ***ε***_*k*_ with E(***ε***_*k*_) = **0** and Cov(***ε***_*k*_) = **I**. For small effects, the GWAS *z*-statistics are 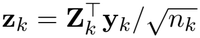. Treating the genotypes as fixed,

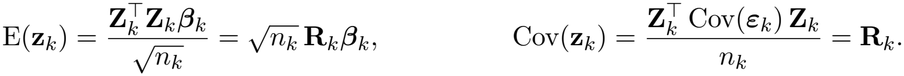

The studies are independent, so Cov(**z**_*k*_, **z**_*l*_) = **0** for *k*≠ *l*, and every cross term of the covariance of the weighted sum vanishes:

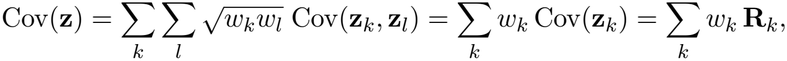

which is Equation (S1). For the mean, *n*_*k*_ = *w*_*k*_*N* gives

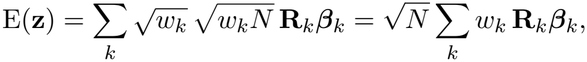

and dividing by 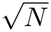 gives Equation (S2). If ***β***_*k*_ = ***β*** for every *k*, the sum is (∑_*k*_ *w*_*k*_**R**_*k*_)***β*** = **R**_mix_***β***. □

##### Assumptions and use in PRS-CS

The result assumes (i) independent studies, since shared participants add cross-study covariance; (ii) the same sample sizes for every SNP, since other-wise the weights become SNP-specific; and (iii) for the mean, effects shared across ancestries on the standardized genotype scale, which the simulations (Section 2.3) satisfy by construction. A GWAS analysed as one pooled sample instead has the LD of a pooled genotype distribution, with a between-ancestry term, as in DISTMIX (Lee et al., 2015), so for pooled designs **D**_mix_ is an approximation. This is also why the two levels of PRS-LDmix use different rules: within an ancestry, its subpopulations are pooled with a between-group term (Section 2.1), because a GWAS analyses them as one sample; across ancestries, the GWAS are meta-analysed, so the LD is the average above, with no between-group term. PRS-LDmix estimates each **R**_*k*_ by a reference panel **D**_*k*_, so **D**_mix_ estimates **R**_mix_, and because the rule needs only correlation matrices, finished PRS-CS panels can be mixed without allele frequencies. In PRS-CS, **D**_mix_ replaces the LD matrix **D** in the likelihood 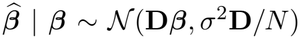 (Ge et al., 2019) and in every sampler update; the prior and sampler are unchanged.

#### S2 SNP alignment across ancestry panels

Mixing adds panels’ LD matrices entry by entry, so for each block every panel must contribute the same SNPs, in the same order, coded with the same effect allele. PRS-LDmix ensures this in four steps.

1. **Common blocks**. All panels use the same block boundaries (Section 2.1), so block *b* covers the same genomic interval in every panel and blocks are matched by index.
2. **Common SNP set**. Per-ancestry filtering on allele frequency can keep different SNPs for different ancestries. Before building the analysis SNP list, the PRS-CS driver intersects the SNP lists of every panel; within each block, the mixing step then keeps only the SNPs present in every panel and in the GWAS summary statistics.
3. **Common order**. Within each block, SNPs are placed in the order of the first panel and every panel’s matrix is reindexed to that order.
4. **Common effect allele**. Allele flips relative to the GWAS effect allele are determined once and the same flips are applied to every panel’s matrix before mixing. This is valid because all panels are built from the same reference index and so code each SNP’s alleles identically. Mixing panels from different sources would require checking alleles panel by panel, which the current scripts do not do.

### Part B: Full simulation results

#### S3 Full simulation results

Tables S1–S4 give the full results of the simulations of Section 2.3, for three ancestry pairs at three compositions; the design and settings are in main-text Tables 1 and 2.

**Table S1:** Top-10% overlap of PRS-LDmix with each single panel, against the same overlap for that panel’s rerun with a new sampler seed. Overlap is the share of target individuals in PRS-LDmix’s top 10% who are also in the panel’s top 10%; the difference is PRS-LDmix minus the seed rerun, in percentage points, and negative values mean PRS-LDmix moves the top 10% further than sampler randomness alone.

| GWAS ancestry mix | Scored in | Top-10% overlap with the panel (%) |  | Difference | Top-10% overlap with the panel (%) |  | Difference |
|---|---|---|---|---|---|---|---|
|  |  | PRS-LDmix | Rerun, new seed | Percentage points | PRS-LDmix | Rerun, new seed | Percentage points |
| **EUR and ASN** |  | Compared with the EUR panel |  |  | Compared with the ASN panel |  |  |
| 90% EUR, 10% ASN | EUR | 91.6 (0.2) | 91.6 (0.2) | 0.0 (0.1) | 76.8 (0.5) | 93.4 (0.2) | −16.6 (0.5) |
|  | ASN | 90.1 (0.2) | 90.2 (0.2) | −0.1 (0.1) | 79.1 (0.3) | 91.0 (0.2) | −11.9 (0.3) |
| 70% EUR, 30% ASN | EUR | 90.6 (0.2) | 91.4 (0.2) | −0.7 (0.1) | 82.7 (0.4) | 92.2 (0.2) | −9.5 (0.5) |
|  | ASN | 89.4 (0.2) | 90.7 (0.2) | −1.3 (0.2) | 84.6 (0.2) | 90.8 (0.2) | −6.3 (0.3) |
| 50% EUR, 50% ASN | EUR | 88.6 (0.2) | 91.1 (0.2) | −2.6 (0.2) | 87.7 (0.2) | 91.4 (0.2) | −3.7 (0.2) |
|  | ASN | 87.2 (0.4) | 91.3 (0.2) | −4.1 (0.4) | 88.3 (0.2) | 90.9 (0.2) | −2.5 (0.2) |
| **EUR and AFR** |  | Compared with the EUR panel |  |  | Compared with the AFR panel |  |  |
| 90% EUR, 10% AFR | EUR | 91.2 (0.2) | 91.1 (0.2) | +0.1 (0.1) | 81.2 (0.3) | 93.9 (0.1) | −12.7 (0.3) |
|  | AFR | 87.4 (0.7) | 87.7 (0.7) | −0.2 (0.6) | 76.0 (1.1) | 91.4 (0.5) | −15.4 (1.0) |
| 70% EUR, 30% AFR | EUR | 88.4 (0.3) | 91.7 (0.2) | −3.3 (0.3) | 82.9 (0.2) | 93.4 (0.1) | −10.4 (0.3) |
|  | AFR | 86.6 (0.8) | 92.4 (0.6) | −5.9 (0.7) | 83.3 (0.9) | 93.1 (0.5) | −9.8 (0.7) |
| 50% EUR, 50% AFR | EUR | 77.7 (0.4) | 94.7 (0.1) | −17.0 (0.5) | 84.2 (0.2) | 94.5 (0.1) | −10.4 (0.2) |
|  | AFR | 82.3 (1.1) | 97.7 (0.3) | −15.4 (1.0) | 86.3 (0.8) | 95.0 (0.4) | −8.6 (0.7) |
| **EUR and SAS** |  | Compared with the EUR panel |  |  | Compared with the SAS panel |  |  |
| 90% EUR, 10% SAS | EUR | 91.6 (0.1) | 91.6 (0.2) | 0.0 (0.1) | 87.1 (0.2) | 92.2 (0.1) | −5.0 (0.2) |
|  | SAS | 89.2 (0.6) | 88.5 (0.6) | +0.7 (0.6) | 84.8 (0.6) | 90.2 (0.7) | −5.5 (0.7) |
| 70% EUR, 30% SAS | EUR | 91.3 (0.1) | 92.2 (0.1) | −0.9 (0.1) | 88.0 (0.2) | 92.7 (0.1) | −4.7 (0.1) |
|  | SAS | 90.3 (0.7) | 90.9 (0.6) | −0.6 (0.6) | 86.0 (0.8) | 90.7 (0.7) | −4.7 (0.7) |
| 50% EUR, 50% SAS | EUR | 88.5 (0.1) | 94.7 (0.1) | −6.3 (0.1) | 88.8 (0.1) | 94.8 (0.1) | −6.0 (0.1) |
|  | SAS | 84.1 (0.8) | 94.7 (0.5) | −10.6 (0.8) | 88.7 (0.7) | 94.8 (0.5) | −6.1 (0.7) |

**Table S2:** As Table S1, for the top 5%, 10% and 20%: PRS-LDmix’s overlap with each single panel minus the seed rerun’s, in percentage points.

| GWAS ancestry mix | Scored in | Overlap difference, PRS-LDmix minus rerun (percentage points) |  |  | Overlap difference, PRS-LDmix minus rerun (percentage points) |  |  |
|---|---|---|---|---|---|---|---|
|  |  | Top 5% | Top 10% | Top 20% | Top 5% | Top 10% | Top 20% |
| **EUR and ASN** |  | Compared with the EUR panel |  |  | Compared with the ASN panel |  |  |
| 90% EUR, 10% ASN | EUR | +0.2 (0.1) | 0.0 (0.1) | +0.1 (0.1) | −19.4 (0.6) | −16.6 (0.5) | −13.2 (0.4) |
|  | ASN | −0.4 (0.2) | −0.1 (0.1) | −0.1 (0.1) | −14.3 (0.4) | −11.9 (0.3) | −9.5 (0.2) |
| 70% EUR, 30% ASN | EUR | −1.1 (0.1) | −0.7 (0.1) | −0.7 (0.1) | −10.6 (0.5) | −9.5 (0.5) | −7.5 (0.4) |
|  | ASN | −1.5 (0.3) | −1.3 (0.2) | −1.0 (0.1) | −7.7 (0.4) | −6.3 (0.3) | −5.3 (0.2) |
| 50% EUR, 50% ASN | EUR | −3.1 (0.2) | −2.6 (0.2) | −2.0 (0.1) | −4.2 (0.3) | −3.7 (0.2) | −2.9 (0.2) |
|  | ASN | −5.7 (0.5) | −4.1 (0.4) | −3.4 (0.3) | −3.1 (0.2) | −2.5 (0.2) | −2.1 (0.1) |
| **EUR and AFR** |  | Compared with the EUR panel |  |  | Compared with the AFR panel |  |  |
| 90% EUR, 10% AFR | EUR | +0.1 (0.2) | +0.1 (0.1) | 0.0 (0.1) | −14.3 (0.4) | −12.7 (0.3) | −9.9 (0.2) |
|  | AFR | −0.5 (0.8) | −0.2 (0.6) | +0.2 (0.3) | −17.2 (1.3) | −15.4 (1.0) | −12.2 (0.6) |
| 70% EUR, 30% AFR | EUR | −4.4 (0.4) | −3.3 (0.3) | −2.7 (0.3) | −12.1 (0.3) | −10.4 (0.3) | −8.4 (0.2) |
|  | AFR | −5.9 (1.0) | −5.9 (0.7) | −4.4 (0.5) | −12.4 (1.0) | −9.8 (0.7) | −7.9 (0.5) |
| 50% EUR, 50% AFR | EUR | −19.9 (0.6) | −17.0 (0.5) | −13.3 (0.4) | −12.5 (0.3) | −10.4 (0.2) | −8.2 (0.2) |
|  | AFR | −19.1 (1.5) | −15.4 (1.0) | −11.5 (0.8) | −11.0 (0.9) | −8.6 (0.7) | −6.1 (0.5) |
| **EUR and SAS** |  | Compared with the EUR panel |  |  | Compared with the SAS panel |  |  |
| 90% EUR, 10% SAS | EUR | 0.0 (0.1) | 0.0 (0.1) | 0.0 (0.1) | −5.9 (0.2) | −5.0 (0.2) | −3.9 (0.1) |
|  | SAS | −2.4 (1.0) | +0.7 (0.6) | 0.0 (0.3) | −6.6 (1.1) | −5.5 (0.7) | −5.0 (0.5) |
| 70% EUR, 30% SAS | EUR | −0.8 (0.1) | −0.9 (0.1) | −0.7 (0.1) | −5.6 (0.2) | −4.7 (0.1) | −3.6 (0.1) |
|  | SAS | −0.6 (1.2) | −0.6 (0.6) | −1.5 (0.4) | −5.6 (1.1) | −4.7 (0.7) | −3.8 (0.5) |
| 50% EUR, 50% SAS | EUR | −7.7 (0.2) | −6.3 (0.1) | −5.1 (0.1) | −7.1 (0.2) | −6.0 (0.1) | −4.9 (0.1) |
|  | SAS | −10.0 (1.2) | −10.6 (0.8) | −7.6 (0.5) | −8.4 (1.2) | −6.1 (0.7) | −4.7 (0.5) |

**Table S3:** As Table S1, for the correlation of scores: the correlation of PRS-LDmix’s scores with each single panel’s, against that of the panel’s seed rerun.

| GWAS ancestry mix | Scored in | Correlation of scores with the panel's |  | Difference | Correlation of scores with the panel's |  | Difference |
|---|---|---|---|---|---|---|---|
|  |  | PRS-LDmix | Rerun, new seed | PRS-LDmix minus rerun | PRS-LDmix | Rerun, new seed | PRS-LDmix minus rerun |
| **EUR and ASN** |  | Compared with the EUR panel |  |  | Compared with the ASN panel |  |  |
| 90% EUR, 10% ASN | EUR | 0.993 (<0.001) | 0.993 (<0.001) | 0.000 (<0.001) | 0.945 (0.002) | 0.995 (<0.001) | −0.050 (0.002) |
|  | ASN | 0.990 (<0.001) | 0.990 (<0.001) | 0.000 (<0.001) | 0.956 (0.001) | 0.992 (<0.001) | −0.036 (0.001) |
| 70% EUR, 30% ASN | EUR | 0.991 (<0.001) | 0.993 (<0.001) | −0.001 (<0.001) | 0.970 (0.001) | 0.994 (<0.001) | −0.024 (0.001) |
|  | ASN | 0.989 (<0.001) | 0.991 (<0.001) | −0.002 (<0.001) | 0.976 (0.001) | 0.992 (<0.001) | −0.016 (0.001) |
| 50% EUR, 50% ASN | EUR | 0.986 (<0.001) | 0.992 (<0.001) | −0.006 (<0.001) | 0.985 (0.001) | 0.992 (<0.001) | −0.008 (0.001) |
|  | ASN | 0.982 (0.001) | 0.992 (<0.001) | −0.010 (0.001) | 0.986 (<0.001) | 0.992 (<0.001) | −0.005 (<0.001) |
| **EUR and AFR** |  | Compared with the EUR panel |  |  | Compared with the AFR panel |  |  |
| 90% EUR, 10% AFR | EUR | 0.992 (<0.001) | 0.992 (<0.001) | 0.000 (<0.001) | 0.964 (0.001) | 0.996 (<0.001) | −0.032 (0.001) |
|  | AFR | 0.986 (0.001) | 0.987 (0.001) | −0.001 (<0.001) | 0.942 (0.002) | 0.992 (<0.001) | −0.050 (0.002) |
| 70% EUR, 30% AFR | EUR | 0.985 (0.001) | 0.993 (<0.001) | −0.008 (0.001) | 0.970 (0.001) | 0.996 (<0.001) | −0.025 (0.001) |
|  | AFR | 0.989 (0.001) | 0.995 (0.001) | −0.006 (<0.001) | 0.974 (0.002) | 0.996 (<0.001) | −0.022 (0.001) |
| 50% EUR, 50% AFR | EUR | 0.943 (0.002) | 0.997 (<0.001) | −0.054 (0.002) | 0.973 (0.001) | 0.997 (<0.001) | −0.024 (0.001) |
|  | AFR | 0.975 (0.003) | 0.999 (<0.001) | −0.024 (0.003) | 0.985 (0.001) | 0.998 (<0.001) | −0.013 (0.001) |
| **EUR and SAS** |  | Compared with the EUR panel |  |  | Compared with the SAS panel |  |  |
| 90% EUR, 10% SAS | EUR | 0.992 (<0.001) | 0.992 (<0.001) | 0.000 (<0.001) | 0.983 (<0.001) | 0.994 (<0.001) | −0.011 (<0.001) |
|  | SAS | 0.990 (<0.001) | 0.990 (<0.001) | 0.000 (<0.001) | 0.978 (0.001) | 0.991 (<0.001) | −0.014 (<0.001) |
| 70% EUR, 30% SAS | EUR | 0.992 (<0.001) | 0.994 (<0.001) | −0.002 (<0.001) | 0.985 (<0.001) | 0.994 (<0.001) | −0.010 (<0.001) |
|  | SAS | 0.990 (<0.001) | 0.992 (<0.001) | −0.003 (<0.001) | 0.982 (<0.001) | 0.993 (<0.001) | −0.011 (<0.001) |
| 50% EUR, 50% SAS | EUR | 0.986 (<0.001) | 0.997 (<0.001) | −0.011 (<0.001) | 0.987 (<0.001) | 0.997 (<0.001) | −0.010 (<0.001) |
|  | SAS | 0.979 (0.001) | 0.998 (<0.001) | −0.018 (0.001) | 0.987 (<0.001) | 0.997 (<0.001) | −0.010 (<0.001) |

**Table S4:**
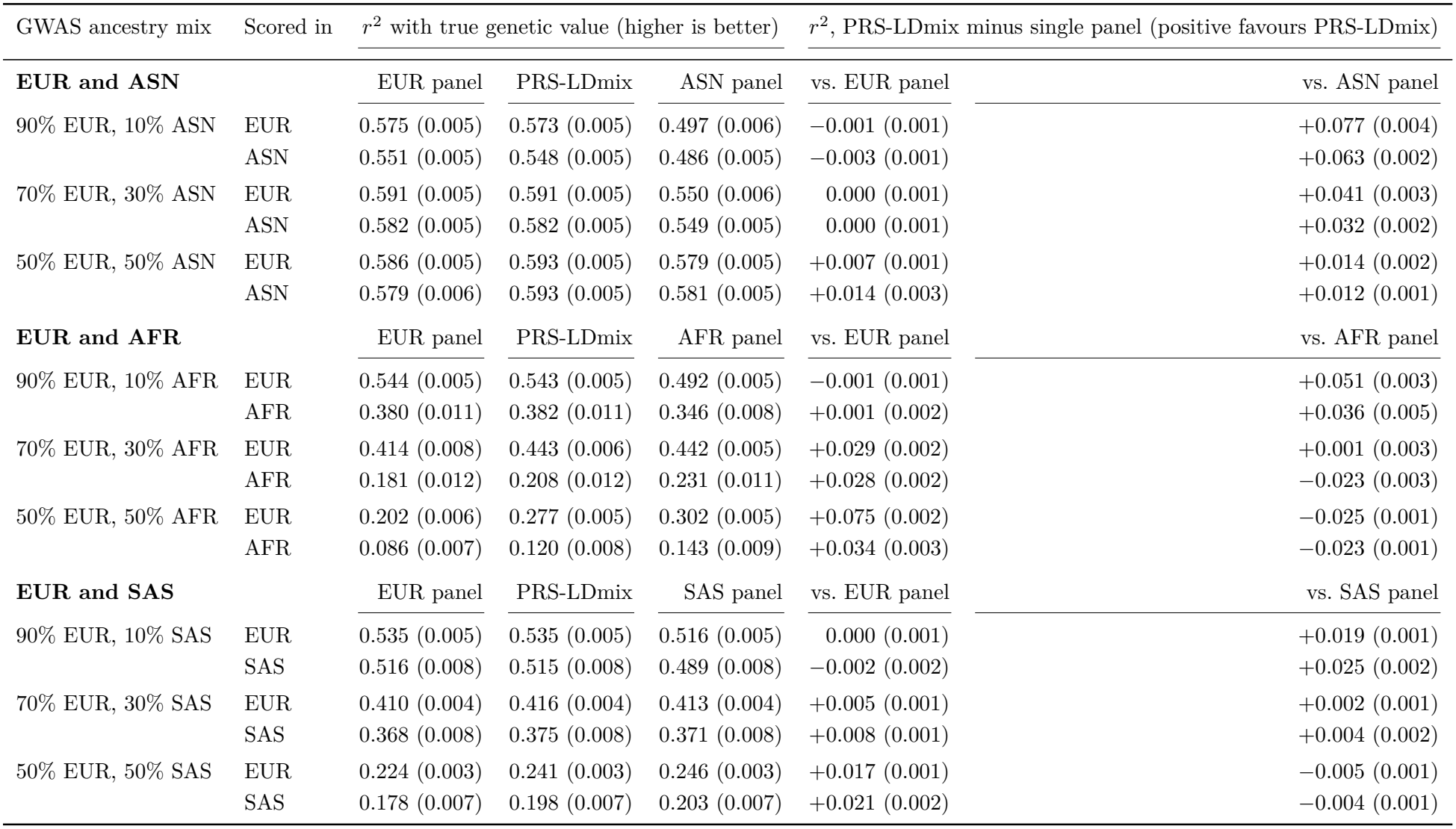
Accuracy of PRS-LDmix and of each single panel: *r*^2^ with the true genetic values, and PRS-LDmix minus each single panel, paired within replicate. Means with Monte Carlo standard errors in parentheses.

| GWAS ancestry mix | Scored in | *r*<sup>2</sup> with true genetic value (higher is better) |  |  | *r*<sup>2</sup>, PRS-LDmix minus single panel (positive favours PRS-LDmix) |  |
|---|---|---|---|---|---|---|
| **EUR and ASN** |  | EUR panel | PRS-LDmix | ASN panel | vs. EUR panel | vs. ASN panel |
| 90% EUR, 10% ASN | EUR | 0.575 (0.005) | 0.573 (0.005) | 0.497 (0.006) | −0.001 (0.001) | +0.077 (0.004) |
|  | ASN | 0.551 (0.005) | 0.548 (0.005) | 0.486 (0.005) | −0.003 (0.001) | +0.063 (0.002) |
| 70% EUR, 30% ASN | EUR | 0.591 (0.005) | 0.591 (0.005) | 0.550 (0.006) | 0.000 (0.001) | +0.041 (0.003) |
|  | ASN | 0.582 (0.005) | 0.582 (0.005) | 0.549 (0.005) | 0.000 (0.001) | +0.032 (0.002) |
| 50% EUR, 50% ASN | EUR | 0.586 (0.005) | 0.593 (0.005) | 0.579 (0.005) | +0.007 (0.001) | +0.014 (0.002) |
|  | ASN | 0.579 (0.006) | 0.593 (0.005) | 0.581 (0.005) | +0.014 (0.003) | +0.012 (0.001) |
| **EUR and AFR** |  | EUR panel | PRS-LDmix | AFR panel | vs. EUR panel | vs. AFR panel |
| 90% EUR, 10% AFR | EUR | 0.544 (0.005) | 0.543 (0.005) | 0.492 (0.005) | −0.001 (0.001) | +0.051 (0.003) |
|  | AFR | 0.380 (0.011) | 0.382 (0.011) | 0.346 (0.008) | +0.001 (0.002) | +0.036 (0.005) |
| 70% EUR, 30% AFR | EUR | 0.414 (0.008) | 0.443 (0.006) | 0.442 (0.005) | +0.029 (0.002) | +0.001 (0.003) |
|  | AFR | 0.181 (0.012) | 0.208 (0.012) | 0.231 (0.011) | +0.028 (0.002) | −0.023 (0.003) |
| 50% EUR, 50% AFR | EUR | 0.202 (0.006) | 0.277 (0.005) | 0.302 (0.005) | +0.075 (0.002) | −0.025 (0.001) |
|  | AFR | 0.086 (0.007) | 0.120 (0.008) | 0.143 (0.009) | +0.034 (0.003) | −0.023 (0.001) |
| **EUR and SAS** |  | EUR panel | PRS-LDmix | SAS panel | vs. EUR panel | vs. SAS panel |
| 90% EUR, 10% SAS | EUR | 0.535 (0.005) | 0.535 (0.005) | 0.516 (0.005) | 0.000 (0.001) | +0.019 (0.001) |
|  | SAS | 0.516 (0.008) | 0.515 (0.008) | 0.489 (0.008) | −0.002 (0.002) | +0.025 (0.002) |
| 70% EUR, 30% SAS | EUR | 0.410 (0.004) | 0.416 (0.004) | 0.413 (0.004) | +0.005 (0.001) | +0.002 (0.001) |
|  | SAS | 0.368 (0.008) | 0.375 (0.008) | 0.371 (0.008) | +0.008 (0.001) | +0.004 (0.002) |
| 50% EUR, 50% SAS | EUR | 0.224 (0.003) | 0.241 (0.003) | 0.246 (0.003) | +0.017 (0.001) | −0.005 (0.001) |
|  | SAS | 0.178 (0.007) | 0.198 (0.007) | 0.203 (0.007) | +0.021 (0.002) | −0.004 (0.001) |

#### S4 LD of the simulated GWAS by ancestry pair

Section 3.2 compares the LD of the simulated GWAS with the LD each analysis assumes. Table S5 summarises that comparison over every block of chromosome 22, and Fig. S1 shows block 75 for EUR+SAS.

**Table S5:**
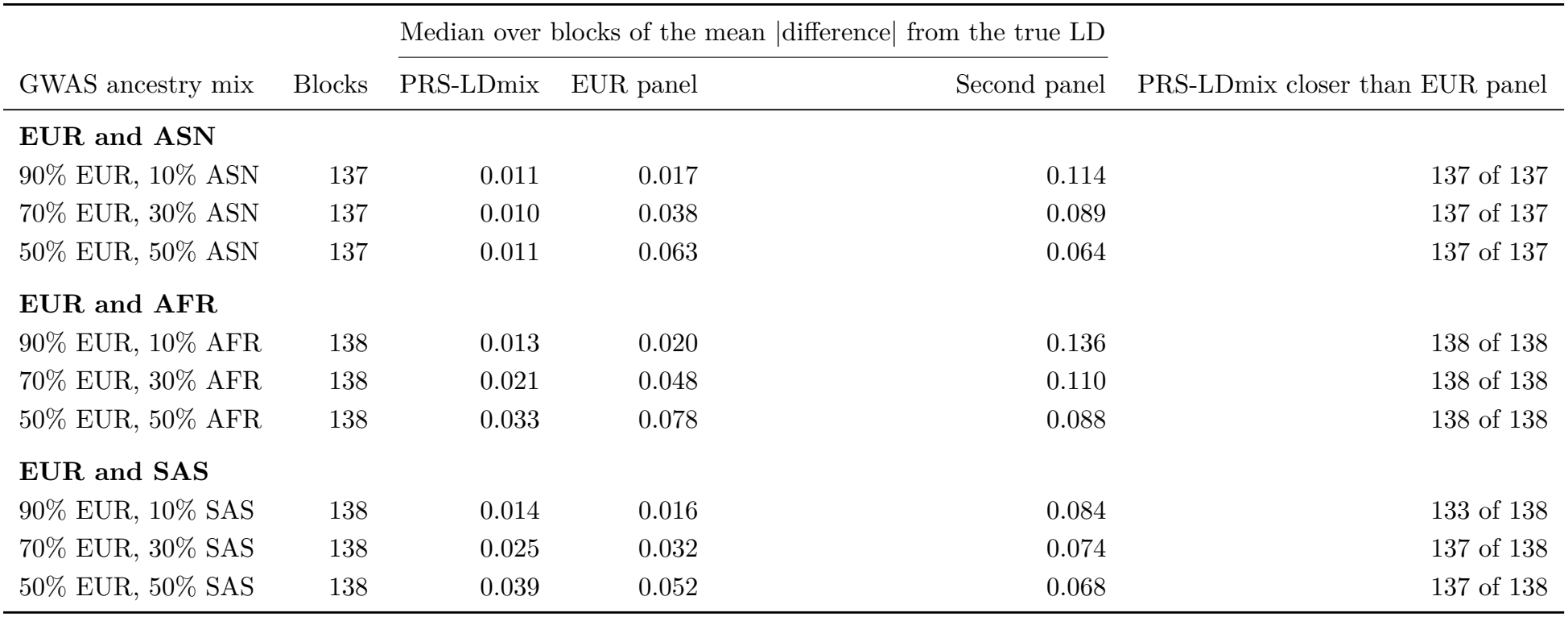
Difference between the true LD of the simulated GWAS and the LD each analysis assumes, over chromosome 22 (fixed 250-kb blocks with at least two analysis SNPs). For each block, the mean |difference| is taken over off-diagonal SNP pairs between the true LD (the discovery LD blended at the GWAS composition) and PRS-LDmix’s blend of the reference panels at the same composition, the EUR panel, or the second ancestry’s panel. Medians over blocks, and the number of blocks where PRS-LDmix is closer to the true LD than the EUR panel. No PRS-CS output and no replicates.

**Figure S1:**
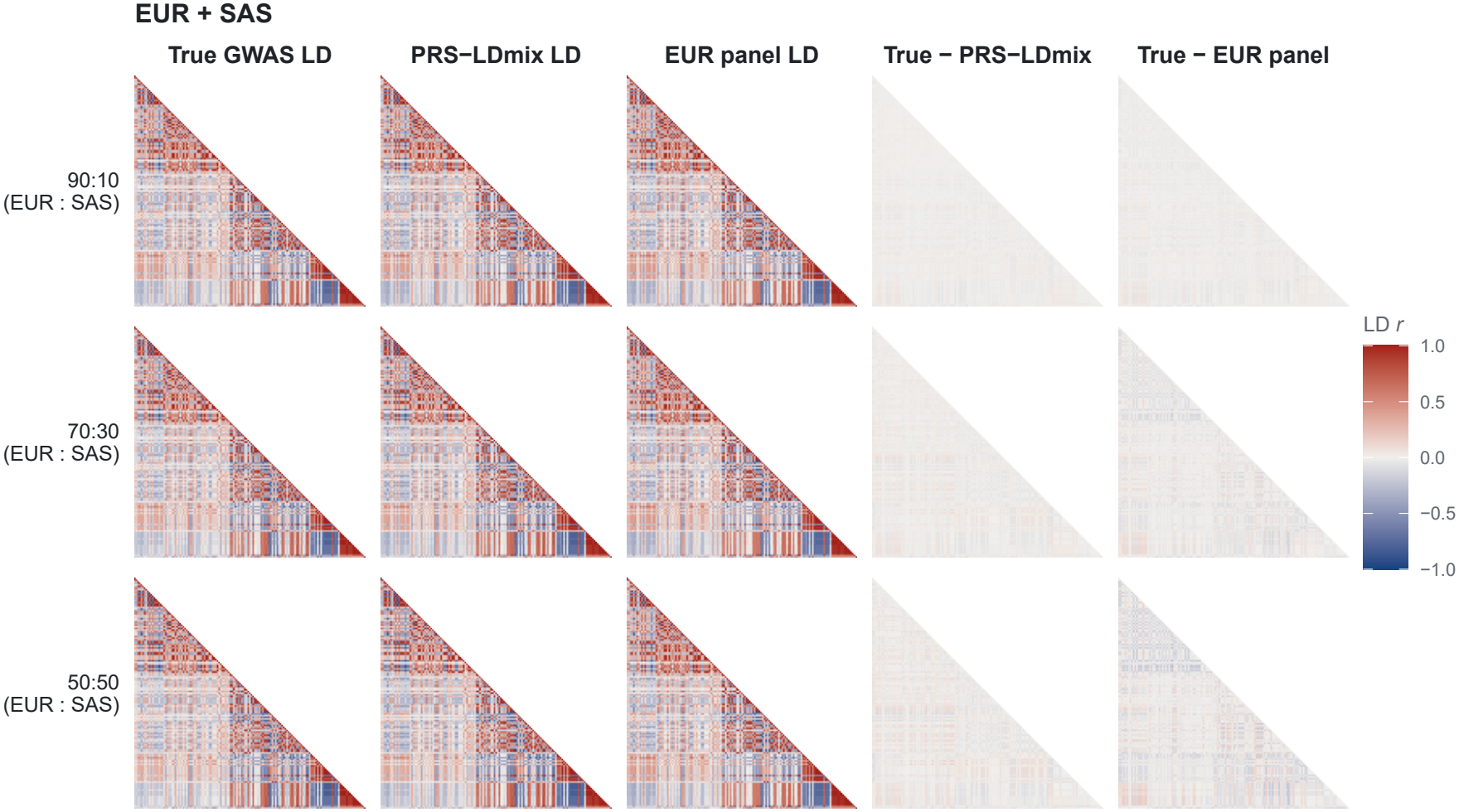
As Fig. 3, for EUR+SAS in the same block; rows are GWAS compositions (EUR : SAS). The SAS discovery cohort is small (132 individuals), so the true LD is noisier and the True ™ PRS-LDmix difference larger (Section 3.2).

### Part C: Robustness checks

#### S5 Panel validation

Genome-wide panels were built from the 33KG reference panel for all five superpopulations, with fixed 250-kb blocks and with EUR-derived cut points, and checked in two ways (Section 3.1): structural and matrix checks, and agreement with PRS-CS’s published 1000 Genomes panels (Table S6); and PRS-LDmix’s script with a single panel against unmodified PRS-CS (Table S7).

**Table S6:**
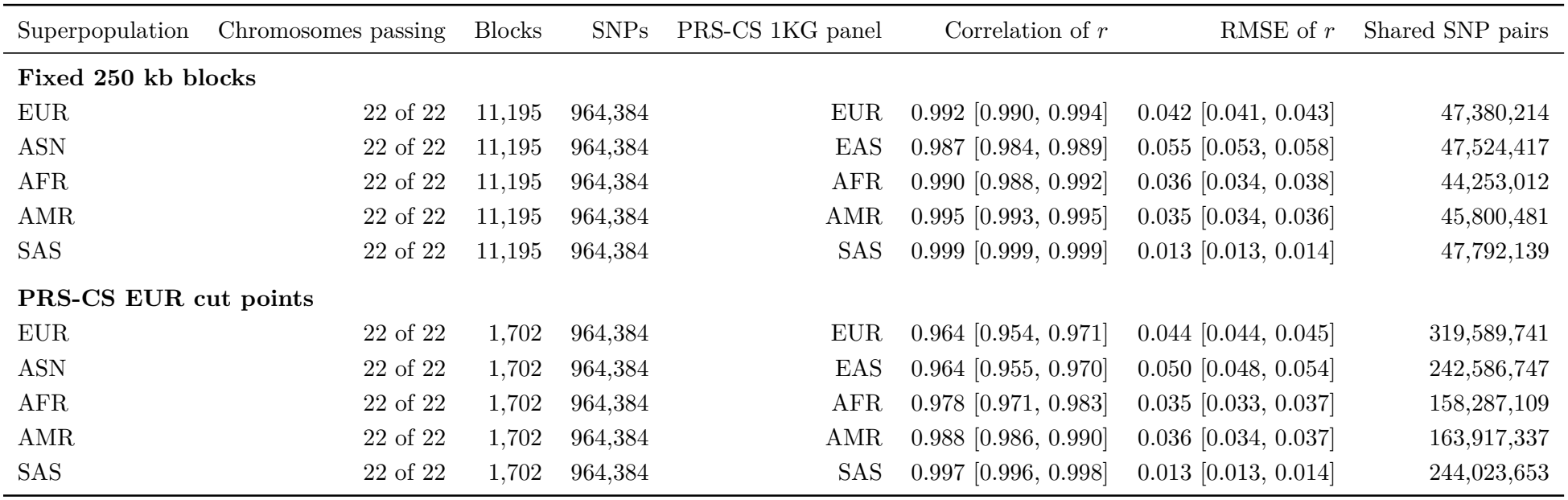
Checks of the genome-wide panels (chromosomes 1–22, HapMap3 SNPs). Chromosomes passing: every panel lists the same SNPs and alleles in the same order, and every block’s LD matrix is finite and symmetric, has a unit diagonal and |*r*| ≤ 1, and is positive definite. Agreement: correlation and root-mean-square difference between our within-block *r* and that of PRS-CS’s published 1000 Genomes panel for the matching population, over SNP pairs in the same block in both panels, after aligning alleles; median and range over the 22 chromosomes. The 33KG panel includes the 1000 Genomes samples, so the comparison checks that the panels are built correctly, not that they replicate in independent samples.

| Superpopulation | Chromosomes passing | Blocks | SNPs | PRS-CS 1KG panel | Correlation of *r* | RMSE of *r* | Shared SNP pairs |
|---|---|---|---|---|---|---|---|
| **Fixed 250 kb blocks** |  |  |  |  |  |  |  |
| EUR | 22 of 22 | 11,195 | 964,384 | EUR | 0.992 [0.990, 0.994] | 0.042 [0.041, 0.043] | 47,380,214 |
| ASN | 22 of 22 | 11,195 | 964,384 | EAS | 0.987 [0.984, 0.989] | 0.055 [0.053, 0.058] | 47,524,417 |
| AFR | 22 of 22 | 11,195 | 964,384 | AFR | 0.990 [0.988, 0.992] | 0.036 [0.034, 0.038] | 44,253,012 |
| AMR | 22 of 22 | 11,195 | 964,384 | AMR | 0.995 [0.993, 0.995] | 0.035 [0.034, 0.036] | 45,800,481 |
| SAS | 22 of 22 | 11,195 | 964,384 | SAS | 0.999 [0.999, 0.999] | 0.013 [0.013, 0.014] | 47,792,139 |
| **PRS-CS EUR cut points** |  |  |  |  |  |  |  |
| EUR | 22 of 22 | 1,702 | 964,384 | EUR | 0.964 [0.954, 0.971] | 0.044 [0.044, 0.045] | 319,589,741 |
| ASN | 22 of 22 | 1,702 | 964,384 | EAS | 0.964 [0.955, 0.970] | 0.050 [0.048, 0.054] | 242,586,747 |
| AFR | 22 of 22 | 1,702 | 964,384 | AFR | 0.978 [0.971, 0.983] | 0.035 [0.033, 0.037] | 158,287,109 |
| AMR | 22 of 22 | 1,702 | 964,384 | AMR | 0.988 [0.986, 0.990] | 0.036 [0.034, 0.037] | 163,917,337 |
| SAS | 22 of 22 | 1,702 | 964,384 | SAS | 0.997 [0.996, 0.998] | 0.013 [0.013, 0.014] | 244,023,653 |

**Table S7:**
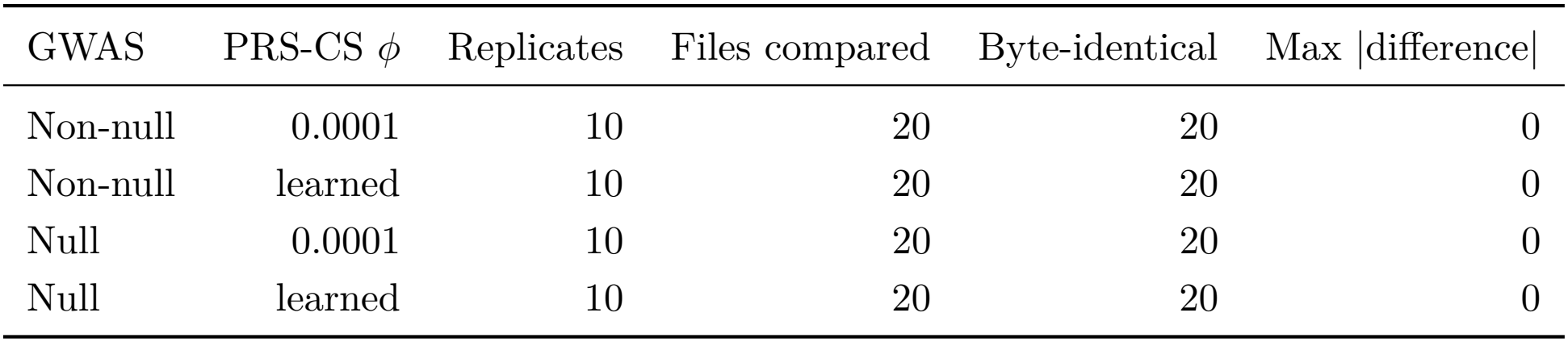
PRS-LDmix’s script (PRScs_mix.py) given a single panel, against unmodified PRS-CS (PRScs.py) with the same panel, GWAS and seed, for both single panels (chromosome 22, 250-kb panels, EUR+ASN 70:30 GWAS, replicates 1–10). Files compared are the posterior SNP-effect files.

#### S6 Other robustness checks

Three further checks ask whether the main results depend on the chromosome, on the reference panels or on the sampler’s chain length.

##### S6.1 A second chromosome

To check that the results are not specific to chromosome 22, two sweep settings were rerun on chromosome 2 with the same individuals: EUR+ASN 70:30 and EUR+AFR 50:50, 100 replicates each, with fixed 250-kb blocks and otherwise the settings of main-text Table 2. Chromosome 2 has several times as many analysis SNPs, so the signal variance *v*_*g*_ was scaled with the SNP count to keep the signal per SNP about the same; EUR+AFR was also run at chromosome 22’s *v*_*g*_, because the AFR discovery sample is small. Changing *v*_*g*_ also changes the signal strength and PRS-CS’s shrinkage, so this comparison does not isolate a single mechanism. Accuracy and score shifts are in Tables S8 and S9.

**Table S8:** *r*^2^ of PRS-LDmix with the true genetic values, and PRS-LDmix minus each single panel (first panel EUR; second ASN or AFR), on chromosomes 22 and 2. Fixed 250-kb blocks, 100 replicates per run; means with Monte Carlo standard errors in parentheses, differences paired within replicate.

| Chromosome, scored in | PRS-LDmix *r*<sup>2</sup> | PRS-LDmix minus first panel | PRS-LDmix minus second panel |
|---|---|---|---|
| **EUR + ASN, 70:30** |  |  |  |
| chr22, scored in EUR | 0.591 (0.005) | 0.000 (0.001) | +0.041 (0.003) |
| chr22, scored in ASN | 0.582 (0.005) | 0.000 (0.001) | +0.032 (0.002) |
| chr2, scaled *v*<sub>*g*</sub>, scored in EUR | 0.358 (0.009) | +0.002 (<0.001) | +0.054 (0.002) |
| chr2, scaled *v*<sub>*g*</sub>, scored in ASN | 0.430 (0.003) | 0.000 (<0.001) | +0.050 (0.001) |
| **EUR + AFR, 50:50** |  |  |  |
| chr22, scored in EUR | 0.277 (0.005) | +0.075 (0.002) | −0.025 (0.001) |
| chr22, scored in AFR | 0.120 (0.008) | +0.034 (0.003) | −0.023 (0.001) |
| chr2, scaled *v*<sub>*g*</sub>, scored in EUR | 0.091 (0.002) | +0.031 (0.001) | −0.023 (0.001) |
| chr2, scaled *v*<sub>*g*</sub>, scored in AFR | 0.077 (0.008) | +0.005 (0.001) | −0.006 (0.001) |
| chr2, *v*<sub>*g*</sub> = 0.01, scored in EUR | 0.079 (0.002) | +0.026 (0.001) | −0.012 (<0.001) |
| chr2, *v*<sub>*g*</sub> = 0.01, scored in AFR | 0.077 (0.008) | +0.006 (0.001) | −0.005 (0.001) |

**Table S9:** PRS-LDmix’s top-10% overlap with each single panel minus the overlap of that panel’s seed rerun, in percentage points, on chromosomes 22 and 2; negative values mean PRS-LDmix moves the top 10% further than sampler randomness alone. Conventions as in Table S8.

| Chromosome, scored in | Compared with the first panel | Compared with the second panel |
|---|---|---|
| **EUR + ASN, 70:30** |  |  |
| chr22, scored in EUR | −0.7 (0.1) | −9.5 (0.5) |
| chr22, scored in ASN | −1.3 (0.2) | −6.3 (0.3) |
| chr2, scaled *v<sub>g</sub>*, scored in EUR | −1.7 (0.1) | −16.3 (0.5) |
| chr2, scaled *v<sub>g</sub>*, scored in ASN | −2.9 (0.2) | −12.8 (0.2) |
| **EUR + AFR, 50:50** |  |  |
| chr22, scored in EUR | −17.0 (0.5) | −10.4 (0.2) |
| chr22, scored in AFR | −15.4 (1.0) | −8.6 (0.7) |
| chr2, scaled *v<sub>g</sub>*, scored in EUR | −19.2 (0.4) | −16.1 (0.3) |
| chr2, scaled *v<sub>g</sub>*, scored in AFR | −11.0 (1.0) | −8.6 (0.7) |
| chr2, *v<sub>g</sub>* = 0.01, scored in EUR | −19.5 (0.4) | −9.6 (0.2) |
| chr2, *v<sub>g</sub>* = 0.01, scored in AFR | −12.0 (1.0) | −5.1 (0.5) |

##### S6.2 Panels built from the discovery individuals

In the main simulations, panels are built from the reference individuals, whose LD differs from that of the discovery individuals the GWAS is simulated on. To test whether this mismatch explains the accuracy of the second-ancestry panel at 50:50 (Section 3.4), the layout pilot also built every panel from the discovery individuals themselves, so that panels differ from the LD the GWAS carries only by the panel estimator (Table S10). These panels are a diagnostic, not a usable method, since a real GWAS’s individual-level genotypes are rarely available.

**Table S10:**
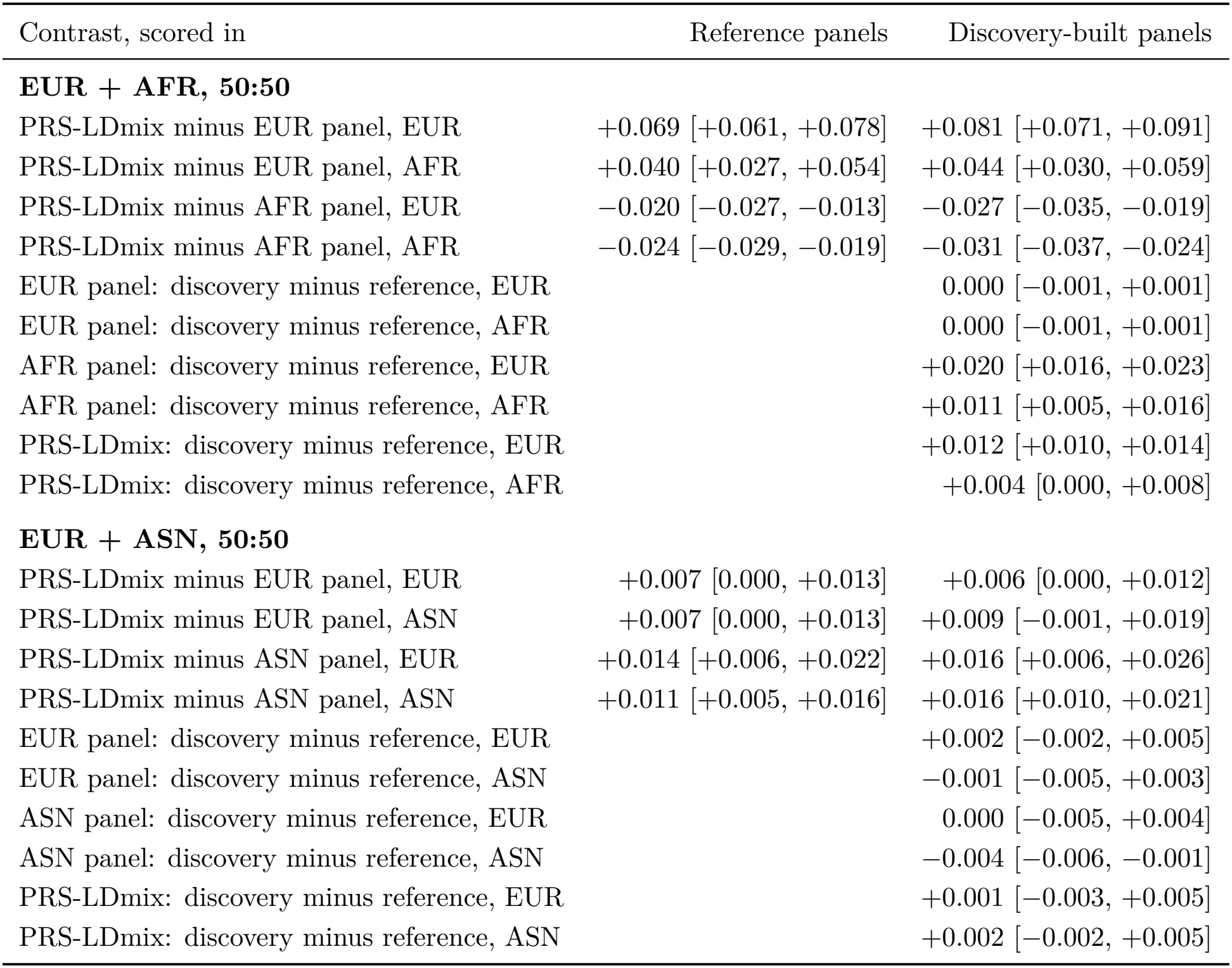
Accuracy with reference panels and with panels built from the discovery individuals. Paired differences in *r*^2^ with the true genetic values, with 95% *t* intervals over 20 replicates (chromosome 22, fixed 250-kb blocks, GWAS composition 50:50). Each contrast is shown with both kinds of panel; the lower rows of each pair give the change in each method’s *r*^2^ from reference to discovery-built panels.

| Contrast, scored in | Reference panels | Discovery-built panels |
|---|---|---|
| **EUR + AFR, 50:50** |  |  |
| PRS-LDmix minus EUR panel, EUR | +0.069 [+0.061, +0.078] | +0.081 [+0.071, +0.091] |
| PRS-LDmix minus EUR panel, AFR | +0.040 [+0.027, +0.054] | +0.044 [+0.030, +0.059] |
| PRS-LDmix minus AFR panel, EUR | −0.020 [−0.027, −0.013] | −0.027 [−0.035, −0.019] |
| PRS-LDmix minus AFR panel, AFR | −0.024 [−0.029, −0.019] | −0.031 [−0.037, −0.024] |
| EUR panel: discovery minus reference, EUR |  | 0.000 [−0.001, +0.001] |
| EUR panel: discovery minus reference, AFR |  | 0.000 [−0.001, +0.001] |
| AFR panel: discovery minus reference, EUR |  | +0.020 [+0.016, +0.023] |
| AFR panel: discovery minus reference, AFR |  | +0.011 [+0.005, +0.016] |
| PRS-LDmix: discovery minus reference, EUR |  | +0.012 [+0.010, +0.014] |
| PRS-LDmix: discovery minus reference, AFR |  | +0.004 [0.000, +0.008] |
| **EUR + ASN, 50:50** |  |  |
| PRS-LDmix minus EUR panel, EUR | +0.007 [0.000, +0.013] | +0.006 [0.000, +0.012] |
| PRS-LDmix minus EUR panel, ASN | +0.007 [0.000, +0.013] | +0.009 [−0.001, +0.019] |
| PRS-LDmix minus ASN panel, EUR | +0.014 [+0.006, +0.022] | +0.016 [+0.006, +0.026] |
| PRS-LDmix minus ASN panel, ASN | +0.011 [+0.005, +0.016] | +0.016 [+0.010, +0.021] |
| EUR panel: discovery minus reference, EUR |  | +0.002 [−0.002, +0.005] |
| EUR panel: discovery minus reference, ASN |  | −0.001 [−0.005, +0.003] |
| ASN panel: discovery minus reference, EUR |  | 0.000 [−0.005, +0.004] |
| ASN panel: discovery minus reference, ASN |  | −0.004 [−0.006, −0.001] |
| PRS-LDmix: discovery minus reference, EUR |  | +0.001 [−0.003, +0.005] |
| PRS-LDmix: discovery minus reference, ASN |  | +0.002 [−0.002, +0.005] |

##### S6.3 Sampler convergence

All analyses use PRS-CS’s default sampler (1,000 iterations, 500 burn-in, thinning by 5). To check that the comparisons between panels do not depend on chain length, we reran EUR+ASN 70:30 with 4,000 iterations (2,000 burn-in) in all 100 replicates (Table S11). Longer chains raised the accuracy of every method by a similar amount, so the differences between PRS-LDmix and the single panels were essentially unchanged.

**Table S11:** Default against longer chains, EUR+ASN 70:30 (chromosome 22, fixed 250-kb blocks, 100 replicates). Upper rows: *r*^2^ with the true genetic values for each method at the defaults and at 4,000 iterations, and the paired change. Lower rows: the paired contrasts between methods at each chain length, and their change. 95% *t* intervals.

|  | *r*<sup>2</sup> at defaults | *r*<sup>2</sup> at 4,000 | Change [95% CI] |
|---|---|---|---|
| *r*<sup>2</sup> **of each arm** |  |  |  |
| EUR panel, scored in EUR | 0.591 | 0.599 | +0.0086 [+0.0067, +0.0105] |
| EUR panel, scored in ASN | 0.582 | 0.590 | +0.0077 [+0.0056, +0.0097] |
| ASN panel, scored in EUR | 0.550 | 0.558 | +0.0077 [+0.0059, +0.0096] |
| ASN panel, scored in ASN | 0.549 | 0.559 | +0.0091 [+0.0075, +0.0108] |
| PRS-LDmix, scored in EUR | 0.591 | 0.599 | +0.0081 [+0.0063, +0.0098] |
| PRS-LDmix, scored in ASN | 0.582 | 0.590 | +0.0082 [+0.0062, +0.0102] |
| **Paired contrasts** |  |  |  |
| PRS-LDmix minus EUR panel, scored in EUR | +0.0003 [−0.0016, +0.0022] | −0.0002 [−0.0017, +0.0012] | −0.0005 [−0.0021, +0.0011] |
| PRS-LDmix minus EUR panel, scored in ASN | −0.0004 [−0.0029, +0.0020] | +0.0001 [−0.0017, +0.0019] | +0.0005 [−0.0015, +0.0026] |
| PRS-LDmix minus ASN panel, scored in EUR | +0.0412 [+0.0356, +0.0469] | +0.0415 [+0.0357, +0.0473] | +0.0003 [−0.0013, +0.0020] |
| PRS-LDmix minus ASN panel, scored in ASN | +0.0323 [+0.0286, +0.0359] | +0.0313 [+0.0275, +0.0352] | −0.0009 [−0.0026, +0.0007] |

#### S7 Block layouts

PRS-CS treats LD as block-diagonal, so any LD between blocks is discarded. The layout pilot compared three layouts on chromosome 22 with the same SNPs: fixed 250-kb tiles (the default), fixed 1-Mb tiles and EUR-derived cut points, inferred from the block structure of PRS-CS’s published EUR panel (Ge et al., 2019). Table S12 reports the change in accuracy when the 250-kb tiles are replaced by either alternative. The interpretation is in the Discussion (Section 4).

**Table S12:** Accuracy by block layout. *r*^2^ with the true genetic values with fixed 250-kb tiles, and the paired change with fixed 1-Mb tiles and with EUR-derived cut points, with 95% *t* intervals over 20 replicates (chromosome 22, reference panels, GWAS composition 50:50). Every layout uses the same SNPs.

| Arm, scored in | *r*<sup>2</sup> at 250 kb | 1 Mb minus 250 kb | EUR cut points minus 250 kb |
|---|---|---|---|
| **EUR + AFR, 50:50** |  |  |  |
| EUR panel, EUR | 0.216 | −0.025 [−0.032, −0.018] | −0.039 [−0.049, −0.029] |
| EUR panel, AFR | 0.090 | −0.004 [−0.008, −0.001] | −0.006 [−0.010, −0.002] |
| AFR panel, EUR | 0.305 | −0.010 [−0.020, 0.000] | −0.018 [−0.030, −0.006] |
| AFR panel, AFR | 0.154 | +0.027 [+0.011, +0.043] | +0.023 [+0.001, +0.045] |
| PRS-LDmix, EUR | 0.286 | +0.002 [−0.008, +0.013] | −0.002 [−0.015, +0.011] |
| PRS-LDmix, AFR | 0.130 | +0.024 [+0.015, +0.033] | +0.026 [+0.014, +0.038] |
| **EUR + ASN, 50:50** |  |  |  |
| EUR panel, EUR | 0.589 | +0.054 [+0.040, +0.068] | +0.083 [+0.068, +0.098] |
| EUR panel, ASN | 0.596 | +0.044 [+0.028, +0.060] | +0.063 [+0.046, +0.079] |
| ASN panel, EUR | 0.582 | +0.043 [+0.028, +0.059] | +0.069 [+0.051, +0.086] |
| ASN panel, ASN | 0.592 | +0.050 [+0.032, +0.068] | +0.080 [+0.061, +0.099] |
| PRS-LDmix, EUR | 0.596 | +0.061 [+0.044, +0.078] | +0.098 [+0.081, +0.115] |
| PRS-LDmix, ASN | 0.602 | +0.062 [+0.042, +0.081] | +0.095 [+0.075, +0.114] |

